# Metabolic priming of GD2 *TRAC*-CAR T cells during manufacturing promotes memory phenotypes while enhancing persistence

**DOI:** 10.1101/2024.01.31.575774

**Authors:** Dan Cappabianca, Dan Pham, Matthew H. Forsberg, Madison Bugel, Anna Tommasi, Anthony Lauer, Jolanta Vidugiriene, Brookelyn Hrdlicka, Alexandria McHale, Quaovi Sodji, Melissa C. Skala, Christian M. Capitini, Krishanu Saha

**Affiliations:** Wisconsin Institute for Discovery, University of Wisconsin-Madison, Madison, Wisconsin USA; Department of Biomedical Engineering, University of Wisconsin-Madison, Madison, Wisconsin USA; Morgridge Institute for Research, University of Wisconsin-Madison, Madison, Wisconsin, USA; Department of Pediatrics, University of Wisconsin School of Medicine and Public Health, Madison, Wisconsin USA; Promega Corporation, Fitchburg, Wisconsin, USA; University of Wisconsin Carbone Cancer Center, University of Wisconsin-Madison, Madison, Wisconsin USA; Department of Human Oncology, University of Wisconsin School of Medicine and Public Health, Madison, Wisconsin USA

**Keywords:** Gene editing, CAR T cells, cancer, metabolism, stem cell memory, CRISPR, GMP-compatible, pre-clinical, scale-up

## Abstract

Manufacturing Chimeric Antigen Receptor (CAR) T cell therapies is complex, with limited understanding of how media composition impact T-cell phenotypes. CRISPR/Cas9 ribonucleoproteins can precisely insert a CAR sequence while disrupting the endogenous T cell receptor alpha constant (*TRAC*) gene resulting in *TRAC*-CAR T cells with an enriched stem cell memory T-cell population, a process that could be further optimized through modifications to the media composition. In this study we generated anti-GD2 *TRAC*-CAR T cells using “metabolic priming” (MP), where the cells were activated in glucose/glutamine low media and then expanded in glucose/glutamine high media. T cell products were evaluated using spectral flow cytometry, metabolic assays, cytokine production, cytotoxicity assays *in vitro* and potency against human GD2+ xenograft neuroblastoma models *in vivo*. Compared to standard *TRAC*-CAR T cells, MP *TRAC*-CAR T cells showed less glycolysis, higher CCR7/CD62L expression, more bound NAD(P)H activity and reduced IFN-γ, IL-2, IP-10, IL-1β, IL-17, and TGFβ production at the end of manufacturing *ex vivo*, with increased central memory CAR T cells and better persistence observed *in vivo*. Metabolic priming with media during CAR T cell biomanufacturing can minimize glycolysis and enrich memory phenotypes *ex vivo*, which could lead to better responses against solid tumors *in vivo*.

## INTRODUCTION

Chimeric antigen receptor (CAR) T cells have emerged as an exciting alternative to traditional cancer treatments for hematologic malignancies with six FDA-approved therapies for multiple myeloma, non-Hodgkin B-cell lymphomas and B cell acute lymphoblastic leukemia available to date. ^1^ In contrast, CAR T therapies for solid tumors have had limited clinical responses, due to a lack of persistence, limited homing to the tumor, and exhausted T-cell phenotypes. ^2, 3^ Clinically, neuroblastoma initially demonstrated the potential efficacy of CAR T cells in treating solid tumors. However, a subsequent analysis revealed a modest 15% response rate with durable responses only achieved in patients whose GD2 CAR T cells persisted longer than 6 weeks after infusion and formed central memory T cells. ^4, 5^ Recently, promising results for advanced neuroblastoma with a third-generation anti-GD2 CAR T cell were reported, with a 63% partial or complete response rate; however, event-free survival remained low at 26%. ^6^ The critical quality attributes that contribute to CAR T cell persistence and central memory formation in solid tumors remain largely unknown.

Manufacturing CAR T-cells to reach therapeutic cell quantities (∼0.1-10 billions cells per patient) involves inducing cell proliferation *ex vivo* through activation with cross-linked antibodies for CD3 and CD28, and cytokine-enriched media.^7–9^ *Ex vivo* culture of CAR T-cells has been previously shown to accelerate differentiation. ^10–12^ Limiting differentiation of these cells into terminal effector or exhausted phenotypes pre-infusion is a goal for the field, since increased stem cell memory fractions in the pre-infusion product have correlated with better cytotoxicity post-infusion, transition to a central memory state, and better persistence *in vivo*. ^13–20^ Enhanced stem cell or central memory phenotypes in CAR T products could be achieved by manipulating media, cytokines, or T-cell metabolism during manufacturing. Tailored media and IL-7/IL-15-based expansion have been found to preserve a stem cell memory profile. ^14^ Metabolic interventions that maintain oxidative phosphorylation (OXPHOS) and suppress glycolysis through metabolic engineering, ^16, 18, 20^ small molecule inhibitors, or glucose/glutamine deprivation ^25–29^ have been shown to enhance the cell persistence, potency, and memory formation of CAR T products. These strategies have yet to be explored with CAR T cell products that have been genome-edited using electroporation (EP) in a virus-free manner.

T cells have classically been transduced with γ-retroviral or lentiviral vectors yielding stable but uncontrolled genomic integration of the CAR transgene with a constitutively active promoter, ^29, 31^ which has been implicated in excessive T-cell differentiation.^33^ CRISPR/Cas9 is an alternative approach that makes a double-stranded DNA break at a precise genomic locus where the DNA repair pathways can stably integrate the desired CAR transgene.^33–35^ This genome editing strategy has been used to insert a CAR transgene upstream of the endogenous T-cell receptor alpha constant (*TRAC*) gene using homology-directed repair (HDR).^36–38^ For current *TRAC* integration strategies, the CAR T cell products lack an intact T cell receptor (TCR) because of the precise genetic knockout of the TCR alpha chain. Relative to conventional viral CAR T products, these *TRAC*-CAR products have more controlled transgene copy numbers in the genome (1 or 2) with CAR transcription driven by the *TRAC* promoter, limited off-target effects, and higher fractions of stem-cell memory phenotypes. ^32, 34^ A benchtop-scale CAR T process for inserting an anti-GD2 construct into the *TRAC* locus using EP of CRISPR/Cas9 ribonucleoproteins with PCR-based donor templates showed promising results in a GD2^+^ human neuroblastoma xenograft model. ^36^ Many studies have optimized culture conditions for CAR T cells manufactured using viral vectors^14, 21, 37^, but it is unknown whether these culture changes will affect *TRAC*-CAR T cells generated by EP in a similar manner.

In this study, we evaluated various culture conditions to develop a flexible, process using GMP-compatible reagents for producing TRAC-CAR T cells at scales suitable for clinical use. We focused on metabolic priming (MP), where T-cells are activated in TexMACS media and then switched post-EP to Immunocult XF. TexMACS media is associated with attenuated cellular activation, possibly due to its lower glucose/glutamine content relative to Immunocult XF.^37, 42^ The result is that MP *TRAC*-CAR T cells are less reliant on glycolysis, exhibit more stem cell memory phenotypes, and consume less glucose than their non-primed counterparts. Furthermore, these MP *TRAC*-CAR T cells can adopt a central memory phenotype following exposure to solid tumors in neuroblastoma xenograft models. This biomanufacturing approach thus holds promise for generating cells with enhanced potency and longevity *in vivo*.

## RESULTS

### *TRAC*-CAR T cell manufacturing at scale

CAR T cells ideally should proliferate *ex vivo* to achieve clinically relevant yields, while maintaining a naïve, stem cell memory/central memory state to achieve persistence *in vivo.* ^10, 15, 43^ We first explored whether a nanoplasmid template could provide a facile method to scale-up production of the donor template for HDR to generate *TRAC*-CAR T cells **(Fig. 1A)**. We electroporated this template along with a CRISPR-Cas9 ribonucleoprotein (RNP) specific for the *TRAC* locus and cultured the cells for 7 days. Robust gene editing at clinical or benchtop scale was observed by flow cytometry, as greater than 90% of T cells lacked expression of the endogenous TCR given the CRISPR-mediated knockout of the *TRAC* gene (**Fig. 1B**) when compared to untransfected T-cells (Clinical: 98.2% (0.7) or Benchtop: 90.7% (1.0) versus Untransfected: 8.6% (5.1), *p* < 0.001 respectively). *TRAC*-CAR T cells were produced equally well at both scales with over 17% CAR positivity using the Lonza or Xenon EP systems (Clinical: 17.1% (0.4) or Benchtop: 22.2% (7) versus Untransfected: 0.02% (0.01), *p* = 0.024 and *p* = 0.049 respectively) **(Fig. 1C)**.

**Figure 1.**
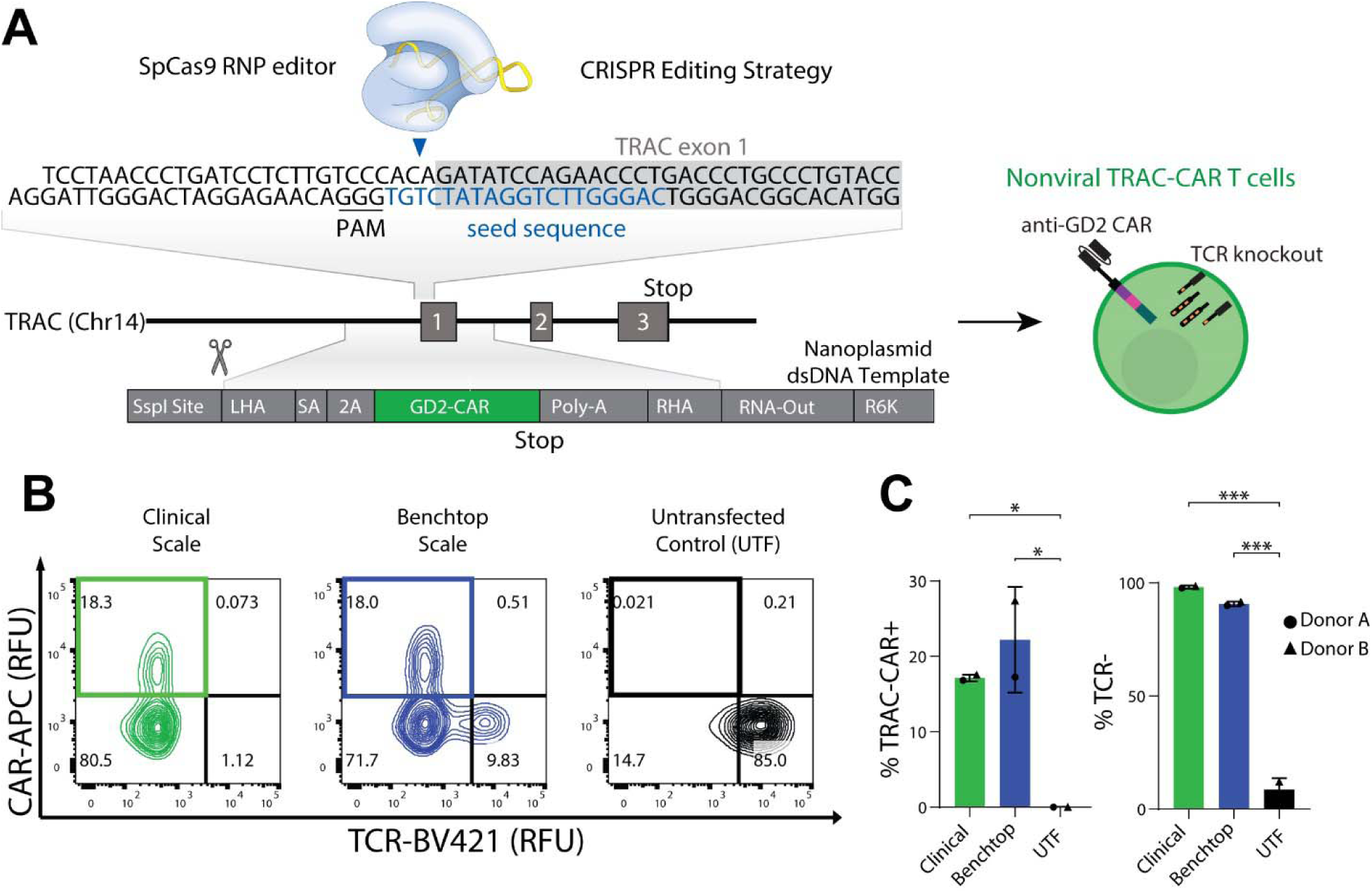
Manufacturing *TRAC*-CAR T cells at clinically relevant scales. (**A**) Schematic of non-viral CRISPR/Cas9 knock-in strategy with a nanoplasmid encoding the CAR. The CAR-containing nanoplasmid encodes for an anti-GD2 CAR under the control of the endogenous *TRAC* promoter. The CAR-containing nanoplasmid has an SSPI restriction site for linearization. (**B**) Representative flow cytometry plots of the relative GD2-CAR vs TCR expression of *TRAC*-CAR T Cells electroporated with SSPI-linearized nanoplasmid on the ThermoFisher Xenon or Lonza 4-D Nucleofector system and untransfected T-cells (Donor A shown). (**C**) Bar graphs comparing GD2 knock- in rates and TCR^-^ percentages of *TRAC*-CAR T cells electroporated on the Xenon or Lonza systems and untransfected T-cells. 2 donors, N_Clinical_=N_Benchtop_=N_Untransfected_ = 2. SA, splice acceptor; 2A, cleavage peptide; LHA, left homology arm; RHA, right homology arm; Poly-A, rabbit β-globin polyA terminator; virus-free CRISPR; CAR, chimeric antigen receptor; *TRAC*, T-cell receptor alpha constant gene; RNP, ribonucleoprotein; PAM, protospacer adjacent motif; TCR, T-cell receptor. Error bars represent mean and standard deviation. Statistical significance was determined with a one-way ANOVA using Dunnett’s T3 test for multiple comparisons (C); *p<0.05; ***p<0.001

### Metabolic priming to enrich for stem cell memory

Generally, CAR T cell production uses a singular media source and cytokine cocktail throughout manufacturing to expand T cells into clinically relevant numbers *ex vivo*. Activation and expansion of *TRAC*-CAR T cells has previously been performed using Immunocult XF medium supplemented with IL-2,^36^ generating stem cell memory T cells with high expression of CD45RA, CD62L, and CCR7 and CAR knock-in rates between 15-34%. ^13, 44^ We sought to optimize culture conditions that could further increase the proportion of stem cell memory *TRAC*-CAR T-cells by manufacturing cells in two media conditions **(Fig. 2A)** using reagents produced via Good Manufacturing Practice (GMP) at a clinically relevant scale: (*1*) standard Immunocult XF Media ^37^ with IL-2 for 10 days (“Control cells”); or (*2*) a transient metabolic priming phase, where cells were activated in TexMACS with IL-7/IL-15 ^21, 45^ before EP and then expanded in Immunocult XF with IL-7/IL-15 for 7 days post-EP (“MP cells”).

**Figure 2.**
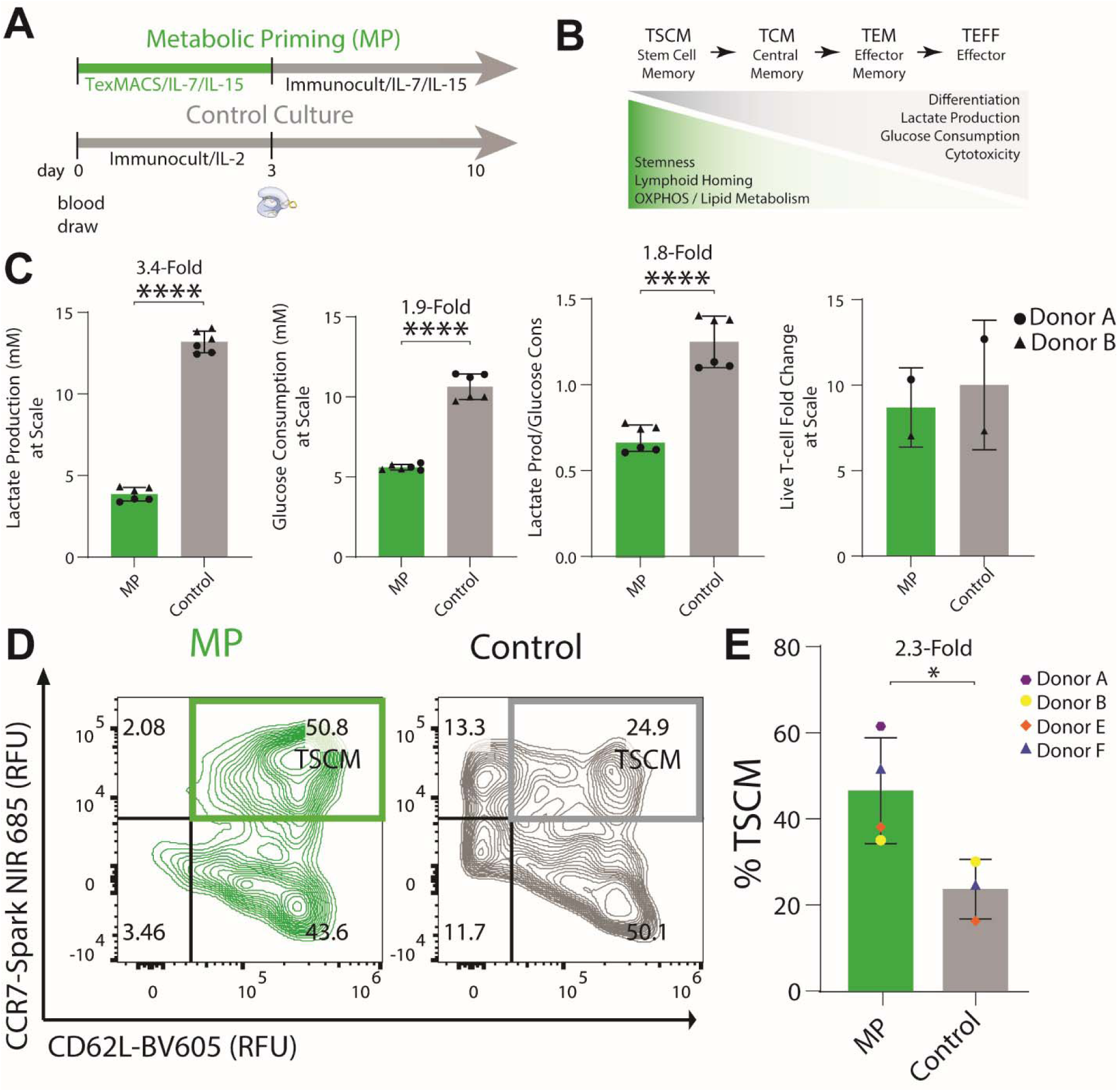
Metabolically primed *TRAC*-CAR T cells produce stem cell memory phenotypes. (**A**) Manufacturing timeline for *TRAC*-CAR T cells: metabolically primed (MP) *TRAC-*CAR T cells are cultured in TexMACS medium for three days during activation and, then Immunocult XF medium supplemented with IL-7/IL-15 for seven days during expansion. Control *TRAC*-CAR T cells are cultured in only Immunocult XF medium supplemented with IL-2 during activation and expansion. **(B)** Schematic of the definition of CAR T cell phenotypes: T_SCM_ (stem cell memory), T_CM_ (central memory), T_EM_ (effector memory), T_EFF_ (effector) as defined by their inherent properties. **(C)** Bar graphs of the lactate production, glucose consumption, lactate production over glucose consumption, and live T-cell fold-change of MP or Control *TRAC*-CAR T cells during expansion in G-Rex6M plates. 2 donors, (lactate/glucose) N_MP_=N_Control_ = 6; (proliferation) N_MP_=N_Control_ = 2 **((D)**Representative contour plots of CD62L/CCR7 co-expression for CAR^+^/TCR^-^ MP and Control *TRAC*-CAR T cells for same conditions. CD62L/CCR7 double positive cells represent a stem cell memory population (T_Scm_) (Donor F shown). **(E)** Bar graph depicting the CD62L/CCR7 double positive population in CAR^+^ T-cells for all conditions. 4 donors, N_MP_ = 4, N_Control_ = 3. Error bars represent mean and standard deviation. Statistical significance was determined with a paired t-test (C), and Welch’s t-test (E); *p<0.05; ****p<0.0001.

Memory T cells primarily use oxidative phosphorylation and fatty acid oxidation for long-term survival, whereas effector T cells rely on glycolysis for rapid proliferation and immediate immune response. In cell cultures, a lower glucose consumption and lactate secretion may indicate a higher proportion of stem cell memory cells (**Fig. 2B**).^16, 18, 20^ We measured the lactate and glucose concentrations of MP and Control *TRAC-*CAR T cells immediately after activation on day 3 and on day 10 of manufacturing to assess their stem cell memory properties. Post-activation MP *TRAC-*CAR T cells produced less lactate (MP: 4.2 mM (2.2) versus Control: 9.2 mM (0.3), *p* = 0.002) and consumed less glucose (MP: 3.6 mM (2.4) versus Control: 13.2 mM (0.5), *p* < 0.001) despite equal fold expansion during activation (MP: 1.1 (0.3) versus Control: 1.2 (0.02)) (**Fig. S1)**. During expansion MP *TRAC*-CAR T cells had produced 1.8-fold lactate per glucose molecule (MP: 0.7 (0.1) versus Control: 1.3 (0.1), *p* < 0.001), 3.4-fold less overall lactate (MP: 3.9 mM (0.4) versus Control: 13.2 mM (0.6), *p* < 0.001) and consumed 1.9-fold less glucose (MP: 5.6 mM (0.2) versus Control: 10.6 mM (0.7), *p* < 0.001) than Control *TRAC-*CAR T cells during expansion in G-Rex6M plates despite having similar yields of cells by Day 10 (MP: 8.7 (1.6) versus Control: 10 (2.7)) (**Fig. 2C**), indicating that metabolic priming does not affect cell yield in this process.

We also demonstrated the importance of transitioning from TexMACs to Immunocult XF with IL-7/IL-15 versus continued culture in TexMACs as the former had the least lactate production and glucose consumption among *TRAC-*CAR T cells manufactured with metabolic priming, TexMACs only, or Immunocult XF only supplemented with IL-7/IL-15 or IL-2 (Fig. S2). MP *TRAC-*CAR T cells also expanded 1.8 times more than those grown in TexMACs only (Fig. S3).

To specifically analyze the phenotypes of the CAR-positive cells in the cell product on day 10, we stained for surface markers of T-cell differentiation and memory phenotypes using a spectral flow cytometry panel and analyzed live CAR^+^/TCR^-^ cells (**Fig. S4).** We classified T cell phenotypes by classic definitions: T_SCM_ (stem cell memory), T_CM_ (central memory), T_EM_ (effector memory) and T_EFF_ (effector). A stem cell memory state is often defined by the expression of surface markers such as CD45RA, CD62L, and CCR7 ^13, 46^ an oxidative versus a glycolytic metabolism, commonly seen in activated effector T-cells, ^22, 47^ and lower glucose consumption. ^28, 48, 49^ (**Fig. 2B**). We defined T_SCM_ *TRAC-*CAR T cells as CCR7^+^/CD62L^+^ which indicates increased capacity for lymphoid homing.^50, 51^ *TRAC*-CAR T cells grown in Immunocult XF are often CD45RA/CD45RO double positive, likely reflecting a transitional state that cannot be assigned to canonical memory subsets.^32, 36^ This transitional phenotype is present in both MP and Control *TRAC-*CAR T cells **(Fig. S5)**. We found that MP *TRAC-*CAR T cells produced 2.3-fold more CD62L^+^/CCR7^+^ CAR^+^ T-cells than Control *TRAC-*CAR T cells (MP: 46.9% (12.3) versus Control: 23.9% (7.0), *p* = 0.028) **(Fig. 2D, E)**. We also observed no significant impacts of priming on CD8 or CD4 expression **(Fig. S5).** *TRAC-*CAR T cells cultured in TexMACs and MP-*TRAC* CAR T cells had similar CD62L/CCR7 expression that did not appear impacted by cytokines (Fig. S2).

### Priming lowers glycolysis and effector phenotypes

To further characterize metabolism, we harvested cells at the on Day 10 of manufacturing for use in a Seahorse assay to analyze oxygen consumption and extracellular acidification rates and for mitochondrial staining for flow cytometry. MP *TRAC-*CAR T cells had lower rates of ECAR (extracellular acidification) and OCR (oxygen consumption) than Control *TRAC-*CAR T cells, indicating lower rates of both glycolysis and overall oxygen consumption **(Fig. 3A)**. Lower basal respiration (MP: 32 pmol/min (14) versus Control: 180 pmol/min (99), *p* < 0.001), maximal respiration (MP: 72 pmol/min (39) versus Control: 298 pmol/min (63), *p* < 0.001), spare respiratory capacity (MP: 40 pmol/min (27) versus Control: 117 pmol/min (56), *p* < 0.001), and basal OCR/ECAR (MP: 0.88 (0.21) versus Control: 1.37 (0.53), *p* = 0.0033) were seen in MP compared to Control *TRAC-*CAR T cells **(Fig. 3B)**. Additionally, MP *TRAC-*CAR T cells had 6% higher normalized mitochondrial mass (MP: 2.3 (0.4) versus Control: 2.2 (0.4), *p* < 0.001, d = 0.33) and 11% higher membrane potential (MP: 1.9 (0.2) versus Control: 1.7 (0.3), *p* < 0.001, d = 0.64) than Control *TRAC-*CAR T cells, further supporting an oxidative phenotype (**Fig. 3C)**. A 27% reduction in granularity (MP: 246 (76) versus Control: 336 (98), *p* < 0.001, d = 1.02) and 14% lower cell size (MP: 310 (52) versus Control: 360 (78), *p* < 0.001, d = 0.74) indicates that MP *TRAC-*CAR T cells may be in a less cytotoxic state^52^ after manufacturing than Control *TRAC-*CAR T cells (**Fig. 3C**).

**Figure 3.**
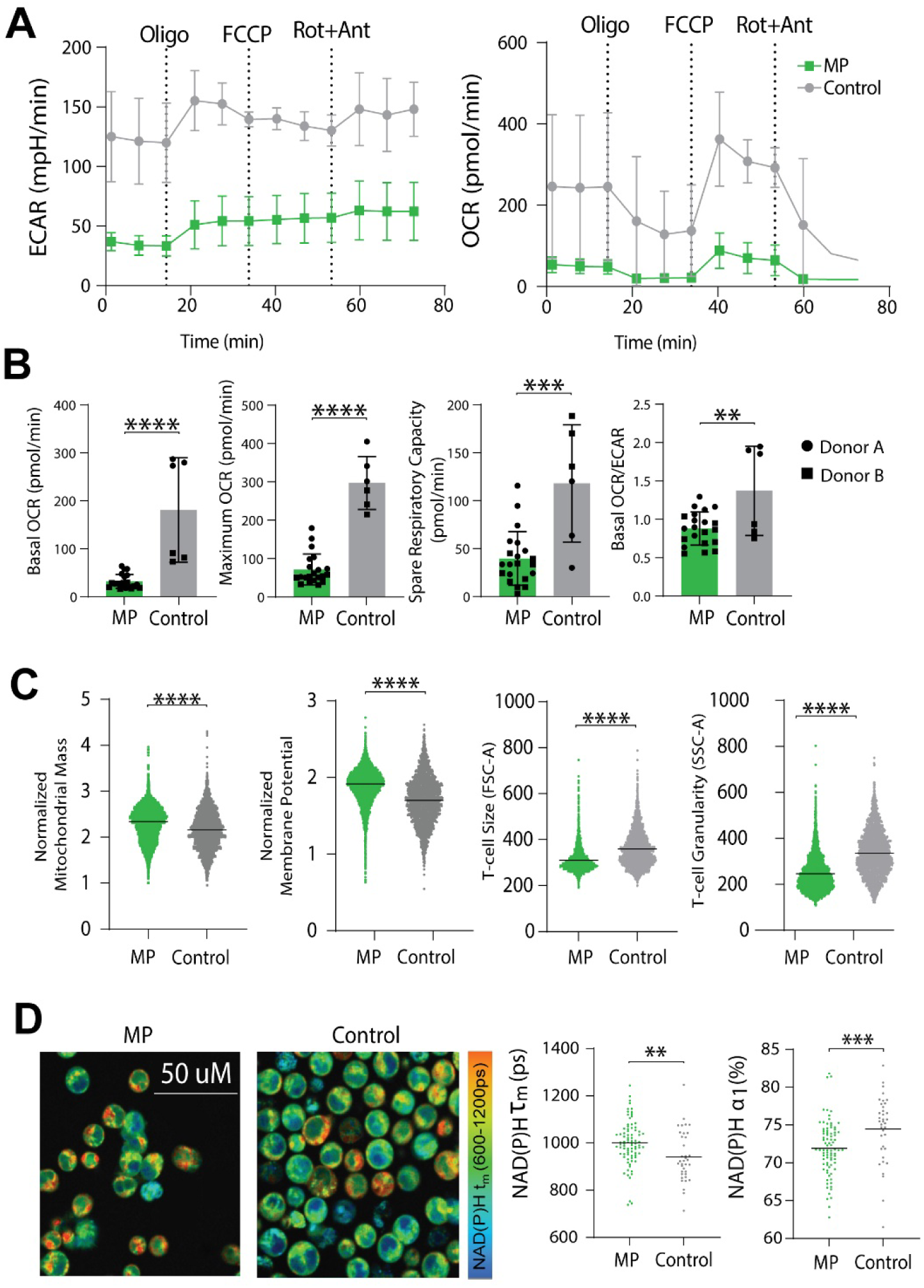
MP *TRAC-*CAR T cells are less glycolytic and have increased mitochondrial mass. **(A)** ECAR and OCR for MP and Control *TRAC-*CAR T cells measured over time by Seahorse (Oligomycin (2.5 *μ*M), FCCP (1 *μ*M), rotenone/antimycin A (0.5 *μ*M)). **(B)** Basal/maximum respiration rates, spare respiratory capacity, and basal OCR/ECAR for MP and Control *TRAC*-CAR T cells measured by Seahorse. 2 donors, N_MP_ = 21, N_Control_ = 6. **(C)** Dot plots for normalized mitochondrial mass (MitoTracker Green intensity divided by FSC-A) normalized mitochondrial membrane potential (TMRE dye intensity divided by FSC-A), cell size (FSC-A), and granularity (SSC-A) for CAR^+^ MP and Control *TRAC*-CAR T cells. 2 donors, N_MP_ = 4264, N_Control_ = 2439. **(D)** Images and bar graphs of NAD(P)H mean lifetime (NAD(P)H τ_m_) and free NAD(P)H fraction (NAD(P)H α_1_) of CAR^+^ MP and Control *TRAC*-CAR T cells as measured by fluorescence lifetime imaging. 2 donors, N_MP_ = 84, N_Control_ = 37 OCR, oxygen consumption rate; ECAR, extracellular acidification rate; SSC-A, side scatter area; FSC-A, forward scatter area; TMRE, tetramethylrhodamine ethyl ester perchlorate. Error bars represent mean and standard deviation. Statistical significance was determined with unpaired t-tests; **p<0.01; ***p<0.001; ****p<0.0001.

To resolve metabolism at single-cell resolution and assess heterogeneity, we also performed fluorescence lifetime imaging of *TRAC-*CAR T cells after manufacturing. We specifically measured autofluorescence lifetimes of NAD(P)H and FAD and report the NAD(P)H mean fluorescence lifetime (NAD(P)H τ_m_) and free NAD(P)H fraction (NAD(P)H α_1_) of CAR^+^ *TRAC-*CAR T cells, which reflect NAD(P)H binding activity. MP *TRAC-*CAR T cells had higher NAD(P)H τ_m_ (MP: 1000 ps (93) versus Control: 942 ps (110), *p* = 0.003) and lower NAD(P)H α_1_ (MP: 72 % (4) versus Control: 75% (4), *p* < 0.001) than Control-*TRAC-*CAR T cells indicating a higher proportion of bound NAD(P)H (**Fig. 3D**) in MP *TRAC*-CAR T cells. These label-free, single cell analyses confirm the presence of more primed cells with increased NAD(P)H binding activity within the MP *TRAC*-CAR T cell products and are consistent with lifetime imaging of similar *TRAC-*CAR products. ^42^

### High potency of MP *TRAC*-CAR T cell products

To assess the potency of the MP *TRAC-*CAR T cells, we measured cytotoxicity and cytokine production after co-culture with the GD2^+^ neuroblastoma cell line, CHLA20. These target cells were seeded at various effector:target (E:T) ratios into 24 or 96-well plates and grown for 24 hours, after which *TRAC-* CAR T cell products were added. The supernatant was taken at 24 hours, and the co-culture was imaged continuously for 48 hours on the Incucyte platform **(Fig. 4A).** MP and Control *TRAC-*CAR T cells achieved similar extents of cytotoxicity for 5:1 and 2.5:1 E:T ratios **(Fig. 4B)**. After 24 hours of co-culture, MP *TRAC-*CAR T cells lysed fewer cancer cells than Control *TRAC*-CAR T cells at 2.5:1 (MP: - 26% (38.1) versus −70.2% (12.1), *p* = 0.041) and 1:1 (MP: 86.5% (38.8) versus Control: −2.5% (29.3), *p* = 0.011) ratios **(Fig. 4C**), but had similar levels of cytotoxicity at the 5:1 ratio. In terms of cytokine production, soluble IFN-γ (MP: 3151 pg/mL (4119) versus Control: 38453 pg/mL (16037), p = 0.0045), IL-2 (MP: 88 pg/mL (121) versus Control: 496 pg/mL (229), *p* = 0.0195), IP-10 (MP: 254 pg/mL (257) versus Control: 17853 pg/mL (8494), p = 0.004), IL-1β (MP: 185 pg/mL (358) versus Control: 16882 pg/mL (14578), *p* = 0.0372), IL-17 (MP: 81 pg/mL (93) versus Control: 664 pg/mL (460), p = 0.0323), and TGFβ (MP: 6.4 pg/mL (13.7) versus Control: 51.9 pg/mL (32.2), *p* = 0.0338) levels produced by MP *TRAC-*CAR T cells in the media were significantly lower than Control *TRAC*-CAR T cells while all other measured cytokines had no differences **(Fig. 4D-F)**. Compared to Control *TRAC*-CAR T cells, MP *TRAC-* CAR T cells have comparable but slower cytotoxic activity with overall lower cytokine production immediately following antigen stimulation *in vitro.* These characteristics have been observed in less differentiated cells, notably T_SCM_ cells. ^13, 49^

**Figure 4.**
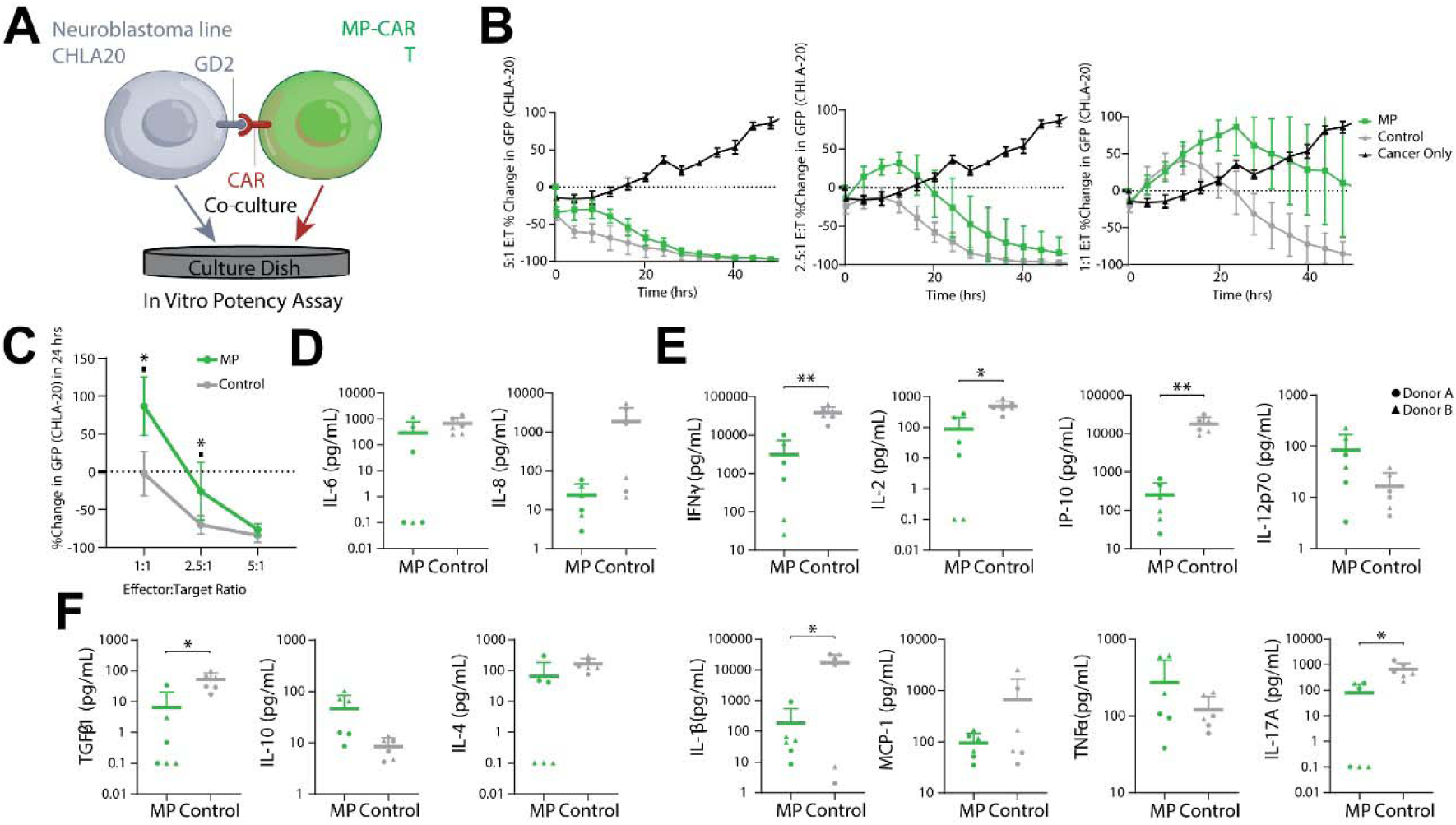
MP *TRAC-*CAR T cells have reduced cytokine production but are equipotent *in vitro* against GD2^+^ neuroblastoma. **(A)** GD2^+^ neuroblastoma CHLA-20 cells were plated in 24 or 96 well plates 24 hrs before *TRAC*-CAR T cell addition. The supernatant was analyzed for cytokine secretion after 24 hours, and potency was measured continuously for up to 48 hours. **(B)** Percentage change in GFP fluorescence from GD2^+^ CHLA-20 neuroblastoma cells versus time in cancer/MP or Control *TRAC*-CAR T cell co-cultures for E:T ratios of 5:1, 2.5:1, or 1:1. **(C)** The percentage change in GFP signal at 24 hours versus E:T ratio is shown for MP or Control *TRAC*-CAR T cells. Co-culture supernatant was analyzed using the LegendPlex Human Essential Immune Response (BioLegend) assay. **(D)** pleiotropic, **(E)** Pro-inflammatory, and **(F)** anti-inflammatory cytokines. The histograms depict the relative levels of secreted cytokines normalized to live T-cell count after co-culture on log scale (pg/mL per 1e6 T-cells). GFP, green fluorescent protein. 2 donors, N_MP_ = 6, N_Control_ = 6. Error bars represent mean and standard deviation. Statistical significance was determined with paired t-tests; *p<0.05; **p<0.01.

To assess the impact of these critical quality attributes on potency *in vivo*, *TRAC-*CAR T cells of both groups were infused into a xenograft NSG mouse model of neuroblastoma. There was similar tumor regression across the two groups **(Fig. S6)**, indicating comparable *in vivo* potency of cells from both culture conditions. Despite lower levels of initial cytotoxicity and cytokine release *in vitro*, MP *TRAC-* CAR T cells demonstrate high potency against solid tumors *in vivo.* Because they were less active in culture, it was unclear if these attributes would make them less prone to exhaustion and exhibit better persistence and central memory formation *in vivo*.

### Enhanced central memory and persistence *in vivo*

To assess memory phenotypes and persistence of MP *TRAC-*CAR T cells *in vivo*, we injected T cell products into a xenograft NSG mouse model for neuroblastoma **(Fig 5A).** Groups included MP *TRAC***-** CAR or Control *TRAC-*CAR cells and No CAR Control cells, which are *TRAC-*T cells expressing an mCherry reporter gene instead of CAR and cultured in the standard “Control” conditions. On day 19 post-injection, we isolated splenocytes, and stained samples for spectral flow cytometry of memory, activation, and exhaustion markers (**Fig. S7**). We found significantly higher amounts of human CD5/CD45^+^ (MP: 2025 cells (3433), Control: 244 cells (487), No CAR Ctrl: 2055 (2841), *p* (MP vs Control) = 0.0423) amounts of lymphocytes in spleens from mice injected with MP *TRAC-*CAR T cells than Control *TRAC-* CAR T cells **(Fig 5B),** indicating improved T cell persistence *in vivo* for MP products. Among transgene^+^/TCR^-^ cells we found no significant differences in CD8 or CD4 expression with nearly 90% of cells being cytotoxic T lymphocytes across conditions (**Fig. S8**).

**Figure 5.**
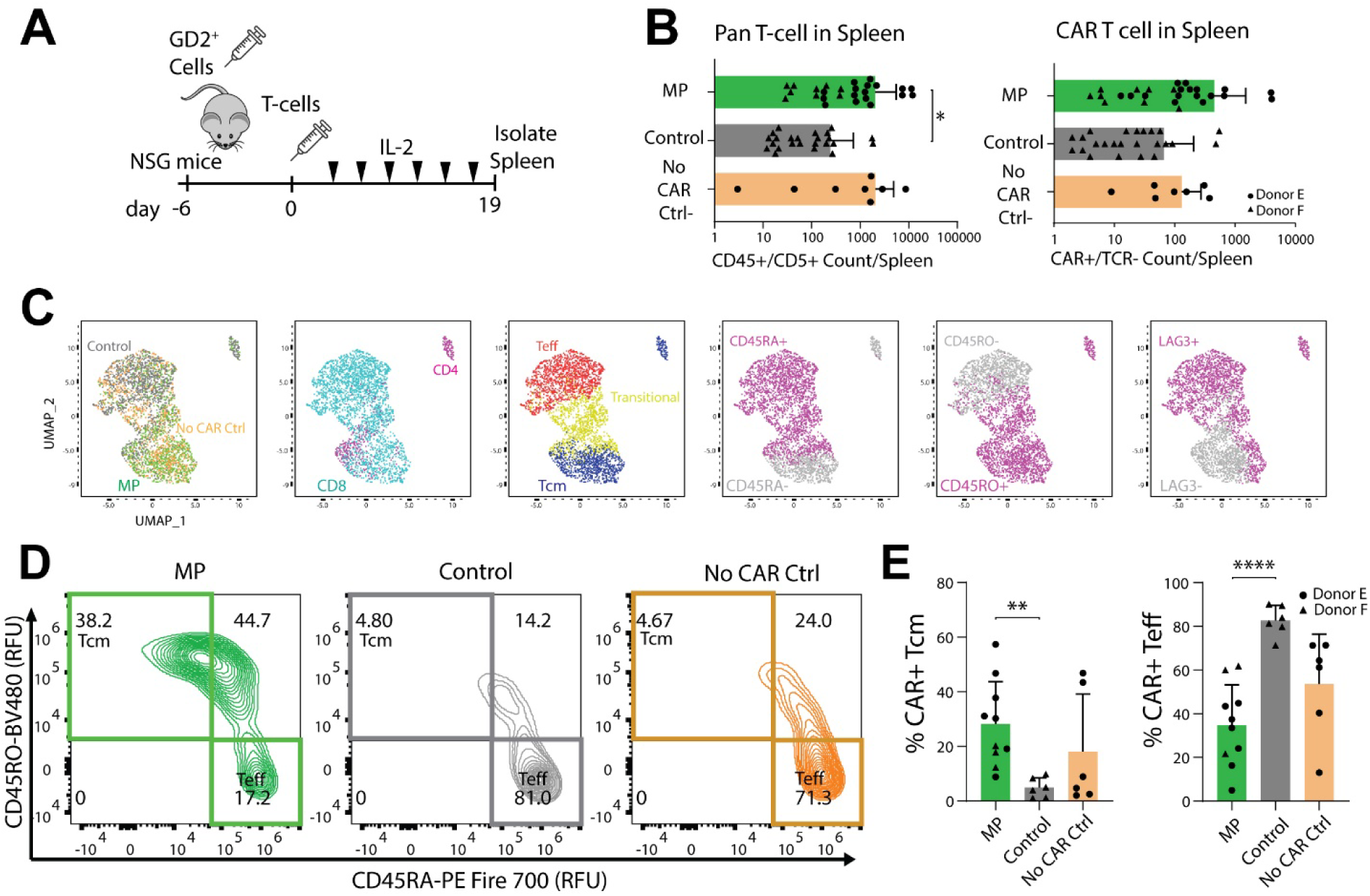
MP *TRAC-*CAR T cells display a central memory phenotype and better persistence *in vivo* after treatment of GD2+ neuroblastoma. **(A)** NSG^TM^ mice were injected with CHLA-20 cells 1 week before *TRAC-* CAR T injection upon which mice were treated with 3 million CAR^+^ cells by tail vein injection. IVIS imaging was performed every 3-4 days with IL-2 supplementation by tail vein. Mouse spleens were isolated on Day 19 post-injection and stained to immunophenotype human T lymphocyte via flow cytometry. **(B)** Bar graphs depicting the number of CD5/CD45^+^ events from isolated mouse spleens (left) and CAR^+^ or mCherry^+^/TCR^-^ lymphocytes within that population for mice treated with MP *TRAC-*CAR T, Control *TRAC*-CAR T, or no CAR Control (mCherry) cells (right) on log scale. 2 donors, N_MP_ = 26, N_Control_ = 24, N_NoCARCtrl_ = 8. **(C)** Marker expression UMAPs of MP, Control *TRAC*-CAR T cells, or No CAR Control T-cells. These maps were generated using flow cytometry to track CD45RA, CD45RO, CD62L, CCR7, LAG3, and TIGIT levels. Dot plots separate cells by condition, CD4/CD8 levels, memory or effector status (CD45RA^-^/CD45RO^+^ (Central memory (T_cm_)), CD45RA^+^/CD45RO^-^ (Effector (T_eff_)), or CD45RA^+^/CD45RO^+^(transitional T-cells)), CD45RO, CD45RA, or LAG3 levels. **(D)** Representative contour plots of the levels of CD45RA versus CD45RO separate MP *TRAC-*CAR T (Donor E), Control *TRAC*-CAR T (Donor F), and no CAR Control cells (Donor E) by central memory or effector status. **(E)** Bar graphs show the relative expression of cell populations for mice with greater than 20 transgene^+^/TCR^-^ events in the spleen: T_cm_ or T_eff_. 2 donors, N_MP_ = 10, N_Control_ = 6, N_NoCARCtrl_ = 6. Error bars represent mean and standard deviation. Statistical significance was determined with Brown-Forsythe and Welch ANOVA tests using Dunnett’s T3 test for multiple comparisons; *p<0.05; **p<0.01; ****p<0.0001.

We assessed splenic lymphocytes according to T cell differentiation and exhaustion using six markers: CD45RA, CD45RO, CD62L, CCR7, LAG3, and TIGIT. MP or Control *TRAC-*CAR T cells separated from each other in a reduced two-dimensional Uniform Manifold Approximation and Projection (UMAP) space (**Fig. 5C**) while no CAR Control cells spanned the entire space more uniformly. The vertical UMAP2 axis separated lymphocytes by differentiation status, with central memory (T_CM_) (CD45RO^+^/CD45RA^-^) and effector (T_EFF_) (CD45RA^+^/CD45RO^-^) T-cells on opposing ends of the axis (**Fig. 5C-D**). This differentiation *in vivo* is expected as T_SCM_ will differentiate after antigen exposure ^53, 54^. T_CM_ cells showed equal CD62L, less LAG3 and CCR7 but more TIGIT expression than T_EFF_ **(Fig. S9)**. Higher percentages of T_CM_ were found in the spleens of mice treated with MP *TRAC-*CAR T cells than with Control *TRAC*-CAR T cells (MP: 28% (16), Control: 5% (4), No CAR Ctrl: 18% (21), *p* (MP vs Control) = 0.0025) (**Fig. 5D, E**), indicating improved central memory MP *TRAC-*CAR T cell persistence *in vivo*. In contrast, there were more T_EFF_ cells in spleens from mice treated with Control *TRAC*-CAR T cells (MP: 35% (18), Control: 83% (7), No CAR Ctrl: 54% (23), *p* (MP vs Control) < 0.001) **(Fig. 5D, E)**.

We also serially stimulated MP or Control *TRAC-*CAR T cells with CHLA-20 cells for 20 days to measure *in vitro* differentiation in response to chronic antigen stimulation. Spectral flow cytometry analysis revealed that MP *TRAC*-CAR T cells had increased persistence of naïve T_SCM_ (MP: 52% (18), Control: 35% (14), *p* = 0.006) and ‘transitional’ (MP: 75% (6), Control: 63% (12), *p* = 0.045) populations relative to Control *TRAC-*CAR T cells. Both groups also separated in a reduced two-dimensional t-SNE space with naïve T-cells clustering closer to MP *TRAC-*CAR T cells (Fig. S10). Overall, these data indicate that MP *TRAC*-CAR T cells show better persistence *in vivo* and *in vitro*, with enrichment of T_CM_ cells noted in the spleen.

## DISCUSSION

We improved upon the manufacturing process of an anti-GD2 *TRAC-*CAR T cell ^36^ by adapting it for higher scales and enriching for desirable stem cell/central memory and metabolic phenotypes. Via priming T-cells in media with lower concentration of glucose, glutamine and potentially other key nutrients, we subsequently reprogrammed T cell metabolism pre- and post-EP during *ex vivo* expansion. Our process used GMP-compatible reagents and can accommodate the EP of 100 million T-cells in a ‘single-shot’ cartridge and up to 25 billion in a ‘multi-shot’ cartridge. ^55^ We used CRISPR/Cas9 to insert a linearized nanoplasmid construct at the *TRAC* locus via HDR, a process that is more scalable than PCR-generated HDR templates due to the facile nanoplasmid amplification within recombinant bacteria. ^56, 57^ Priming T-cells with TexMACS supplemented with IL-7/IL-15 pre-EP to briefly slow glycolysis and proliferation followed by expansion in Immunocult XF with IL-7/IL-15 post-EP produced a final MP *TRAC*-CAR T product enriched for stem cell memory T cells with a unique metabolic profile that showed comparable potency to *TRAC*-CAR T generated under a single media source, but better persistence and higher enrichment of central memory T cells *in vivo*, an ongoing goal of the field of CAR T therapy for solid tumors.

Compared to Control cells, MP *TRAC*-CAR T cells had higher populations of CD62L^+^/CCR7^+^ cells, lower lactate production, glucose consumption, and ECAR post-manufacturing, making them distinctly more T_SCM-_like^13, 16, 49^ (**Fig. 2C, 2F, 3A**). These cells also had lower lactate production per mole of glucose consumed indicating a shift away from glycolysis (Fig 2C). Additionally, MP *TRAC*-CAR T cells had higher mitochondrial membrane potential and mass when normalized to cell size **(Fig. 3B)** which is often seen in memory cells as they switch to OXPHOS and fatty acid oxidation metabolism. ^47^ Surprisingly, these cells also had lower rates of oxygen consumption and a lower OCR/ECAR (**Fig 3A**): this finding is consistent with another study on the effects of intermittent fasting *in vivo* in murine bone-marrow T-cells where a similarly low ECAR and OCR were observed along with reduced mTOR activity. ^58, 59^ The reduced OCR and ECAR in MP *TRAC-*CAR T cells could be due to reduced mTOR signaling, considering that mTOR is upregulated in effector T-cells ^60^ and MP *TRAC-*CAR T cells were less cytotoxic than Control *TRAC*-CAR T s *in vitro* **(Fig. 4),** but further studies of the role of mTOR within MP *TRAC*-CAR T cells are needed.

The lower OCR, ECAR, and glucose consumption in MP *TRAC*-CAR T cells may have also reduced granzyme production, mediators of cytotoxic activity, especially given the lower granularity (side scatter by flow cytometry) in MP *TRAC*-CAR T cells. However, this reduced granularity *ex vivo* is overcome *in vivo* as both culture conditions produced *TRAC-*CAR T cells that can successfully suppress tumor growth in a xenograft mouse model **(Fig. S6)**. An advantage of MP *TRAC*-CAR T cells was the generation of more central memory cells compared to Control *TRAC*-CAR T cells *in vivo* **(Fig. 5),** with better persistence in the periphery (spleen). These cells also maintained a more naïve phenotype in *in vitro* serial stimulation assays, indicating delayed differentiation in MP *TRAC-*CAR T cells when chronically activated. Given this known limitation for retrovirally-transduced anti-GD2 CAR T cells generated with standard biomanufacturing techniques to treat neuroblastoma ^5^, clinical trials with virus-free GD2 *TRAC*-CAR T cells with a MP biomanufacturing process will be needed to determine if enhanced persistence and central memory can be achieved in patients.

Our novel metabolic priming process demonstrates how modulating media composition and cytokines of *TRAC-*CAR T cells pre- and post-EP influences CAR T-cell phenotype, metabolism and persistence post-infusion. While our study used *TRAC-*CAR T cells, there were no significant differences in CD62L/CCR7 expression based on CAR or TCR expression in our product. Additionally, primary T-cells undergoing metabolic priming have similar drops in lactate production and glucose consumption compared to control cells (Fig. S11), suggesting the benefits of metabolic are not dependent on CAR or TCR expression and may extend to other modalities. While many studies have tuned culture conditions of viral CAR T-cells to increase metabolic fitness, T_SCM_ properties, and potency, ^16, 17, 25, 27, 29^ it is unclear if these benefit *TRAC-* CAR T cells, already having been shown to have more potent memory phenotypes. ^41^ We and others have shown the importance of restricting glycolysis only during activation as expanding *TRAC-*CAR T cells in TexMACs ablates the metabolic phenotype adopted by MP *TRAC-*CAR T cells (Fig. S2) and produces a lower cell yield (Fig. S3). ^42^ Activating T-cells in TexMACs rewires their metabolism away from glycolysis (Fig. S1) and it may be possible that factors in Immunocult XF can uniquely enforce this phenotype post-EP. Higher glutamine content in Immunocult-XF compared to TexMACs ^17^ may create a glutamine deprivation during activation, which has been shown to increase metabolic fitness of therapeutic T-cells. ^25, 26^ Lower glutamine during expansion in TexMACs may force T-cells to uptake more glucose to compensate and lower cell yield. Additionally, while the media choice during *TRAC-* CAR T cell culture seems to impact metabolism more than the choice of cytokines, priming anti-GD2 *TRAC-*CAR T cells with IL-2 could produce similar stem cell memory phenotypes with enhanced *in vivo* performance. ^42^ Additional strategies will need to be explored to determine if these effects can be enhanced even further. For example, treating T-cells during activation with N-acetylcysteine, a well-known antioxidant, can promote expression of stem cell memory markers and lower glycolysis. ^28^ T cell metabolism could be manipulated using genetic engineering ^30^, as *PRODH2* overexpression can shift cells to an OXPHOS-based metabolism with more active mitochondria and an increased percentage of CD45RA^+^/CD62L^-^ T-cells post-tumor challenge.^30^ Transient glucose restriction and treatment of T-cells with the glutamine inhibitor DON have also both been shown to increase T_SCM_ surface marker expression and lower glycolysis. Advances to *TRAC-*CAR T cell biomanufacturing has the capacity to produce cell therapies with favorable biologic characteristics that could potentially improve CAR T cell responses against solid tumors.

## MATERIALS AND METHODS

### Cell Lines

GD2^+^ human neuroblastoma CHLA-20 cells were gifted by Dr. Mario Otto (University of Wisconsin-Madison). These cells were cultured in Dulbecco’s Modified Eagle Medium (DMEM) supplemented with 10% fetal bovine serum (FBS) (Gibco, Thermo Fisher Scientific, Waltham, MA) and 1% penicillin-streptomycin (P/S) (Gibco, Thermo Fisher Scientific, Waltham, MA). AkaLucGFP CHLA-20 cells were created through viral transduction by Dr. James Thomson (Morgridge Institute for Research). In short, Phoenix cells (ATCC, Manassas, VA) were grown in DMEM with 10% FBS and 1% P/S and selected with 1 ug/mL of diphtheria toxin (Cayman Biologics, Ann Arbor, MI) and 300 ug/mL hygromycin (ThermoFisher Scientific, Waltham, MA). Selection for transgene-positive cells was confirmed by flow cytometry for mouse Lyt2 expression as a reporter gene (Biolegend, San Diego, CA). 3T3 cells were grown in DMEM with 10% FBS and 1% P/S. Cell authentication was performed using short tandem repeat analysis (Idexx BioAnalytics, Westbrook, Maine, USA) and per ATCC guidelines using cell morphology, growth curves, and *Mycoplasma* testing within 6 months using the MycoStrip Mycoplasma Detection Kit (Invitrogen, Waltham, MA). Cell lines were maintained in culture at 37*°*C in 5% CO_2._

### Plasmid Constructs

A GD2-CAR plasmid construct encoding a 2A.14G2A-CD28-OX40-CD3ζ CAR gifted by Malcolm Brenner (Baylor College of Medicine) was synthesized and the sequence verified (GenScript, Piscataway, NJ). A separate no CAR Control (mCherry) construct was contained in an H2B-mCherry sequence in place of the GD2-CAR and designed, synthesized, and sequenced the same as the GD2-CAR plasmid (Genscript, Piscataway, NJ). Both transgenes were flanked by 500 bp homology arms and cloned into a pUC57 backbone, grown in 5-alpha competent *Escherichia coli* (NEB, Ipswich, MA) and purified using the PureYield MidiPrep System (Promega, Madison, WI).

### Sanger Sequencing

The GD2-CAR and no CAR Control had their sequences verified via Sanger Sequencing at the UW-Biotechnology Center. Briefly, each construct was separated into 20 uL aliquots with appropriate primers for sequencing found in Supplementary Table 1. PCR was performed, the amplicons separated by gel electrophoresis, and peaks analyzed for sequence identity.

### Nanoplasmid Production

Linear dsDNA templates were made via PCR amplification was done using the GD2-CAR or no CAR Control plasmid as a template using Q5 Hot Start Polymerase (Cat # M0494S, NEB, Ipswich, MA) in 50 uL reaction volumes. The cycling parameters were 98°C for 10 s, 65°C for 20 s, and 72°C for 90s, for a total of 35 cycles. These reactions were then pooled into 600 uL reactions for Solid Phase Reversible Immobilization (SPRI) cleanup (1x) using AMPure XP (Cat # B23318, Beckman-Coulter, Brea, CA) beads according to the manufacturer’s instructions and eluted at 2 mg/mL in DNAase free water. The linear products were shipped to Aldevron (Fargo, ND) where they were blunt cloned into a nanoplasmid backbone consisting of two components: R6K origin of replication and an anti-levansucrase RNA sequence that enables antibiotic free selection. The RNA-OUT platform prevents the expression of SacB which prevents toxicity from Levansucrase as well as transgene silencing after genomic insertion. ^62, 63^ Nanoplasmid DNA was manufactured on-site at Aldevron and resuspended in DNAase free water a 2 mg/mL. Nanoplasmid sequences can be found in Supplementary Table 2. Primer sequences are shown in Supplementary Table 3.

### Nanoplasmid Linearization by Restriction Digest

To linearize nanoplasmid for use as a dsDNA HDR template, a restriction digest of the nanoplasmid constructs using SSPI-HF (Cat # R3132S, New England Biolabs, Ipswich, MA) was performed. Four restriction digest batch reactions in 1.5 mL Eppendorf tubes (50 uL nanoplasmid, 125 uL CutSmart buffer, 25 uL SSPI-HF enzyme, and 1050 uL DNAase-free water for 1250 uL total) were aliquoted (50 uL into PCR tubes for a total of 96 reactions). These were incubated in a thermocycler at 37°C for 15-60 min and heat inactivated at 65°C for 20 min according to the manufacturer’s instructions. Gel electrophoresis was then performed on the finished product to assess if proper cutting took place, followed by SPRI cleanups to purify and concentrate the material to 2 mg/mL. PCR reactions were pooled into eight 1.5 mL Eppendorf tubes (600 uL) with an equal volume of SPRI (Beckman-Coulter, Brea, CA) beads that incubated for 5 minutes at room temperature. The product was washed twice with 70% ethanol and eluted in 75 uL of DNase-free water and pooled into one tube (600 uL). This product was subject to a second round of cleanups and was eluted in 30 uL of water. DNA was quantified using the NanoDrop2000 Qubit double-strand DNA (dsDNA) Broad Range (BR) Assay (ThermoFisher Scientific, Waltham, MA) and diluted to 2 mg/mL according to Qubit measurements.

### Isolation of T-cells from Peripheral Blood

Peripheral blood was drawn from healthy donors using an IRB-approved protocol (2018-0103). Blood was collected into Lithium Heparin-coated vacutainer tubes and transferred to 50 mL conical tubes. CD3^+^ primary human T-cells were isolated by negative selection as per manufacturer’s instructions (Cat # 15021, 15061, RosetteSep Human T-cell Enrichment Cocktail, STEMCELL Technologies, Vancouver, Canada). T-cell pellets were resuspended in dilution medium and counted using a hemocytometer with 0.4% Trypan blue viability stain (ThermoFisher Scientific, Waltham, MA). Cells were then resuspended at 1 million/mL in either Immunocult-XF T-cell Expansion Medium (Cat # 10981, STEMCELL Technologies, Vancouver, Canada) or TEXMACs Cell Culture Medium (Cat # 130-097-196, Miltenyi Biotec, Bergisch Gladbach, Germany). T-cell cultures were supplemented with 200 U/mL IL-2 (Cat # 200-02, Peprotech (Thermo Fisher), Cranbury, NJ) or 10 ng/mL of IL-7 (Cat # 207-IL-005/CF, BioTechne, Minneapolis, MN) and 10 ng/mL of IL-15 (Cat # 247-ILB-005/CF, BioTechne, Minneapolis, MN) and stimulated with Immunocult Human CD3/CD28/CD2 T-cell Activator (25 uL for each mL of culture, Cat # 10990, 10970, STEMCELL, Vancouver, Canada) for 48 hours or T-cell TransAct (10 uL for each mL of culture, Cat # 130-111-160, Miltenyi Biotec, Bergisch Gladbach, Germany) for 72 hours respectively.

### Isolation of T-cell from LRS Cone

LRS cones (Versiti Blood Bank, Milwaukee, WI) were purchased as an alternative to drawing from healthy donors. Briefly, RBC-depleted peripheral blood was flushed out of the cone with a blunt gauge needle and dilution medium. The solution was diluted 1:1 with dilution medium and 30 mL was carefully layered on top of 15 mL of Lymphoprep (Cat # 07811, STEMCELL Technologies, Vancouver, Canada) solution. Tubes were centrifuged for 1200 g x 20 min with the brake off. The white monolayer containing leukocytes was then gently disturbed and pipetted into a 15 mL conical tube which was spun at 300 g x 5 min and washed twice. T-cells were then positively selected using an EasySep T-cell kit (Cat # 17951, STEMCELL Technologies, Vancouver, Canada) and an EasySep Magnet (STEMCELL Technologies, Vancouver, Canada) as per the manufacturer’s protocol. T-cell supernatant was collected from the magnet and the cells were then counted, resuspended in culture medium, and activated as above.

### T-cell Electroporation on Lonza 4D Nucleofector

Following T-cell activation, RNP’s and DNA were electroporated (EP) into T-cells in 16 well EP cuvettes on a 4-D Nucleofector X unit (Cat # V4XP-3032, Lonza, Walkersville, VA) using pulse code EH-115. One million cells were electroporated per well in the cuvette. Per reaction, a single guide RNA (IDT, Coralville, IA) specific for the TRAC locus (2 uL of 100 uM, IDT) was incubated with SpCas9 (0.8 uL of 10 mg/mL, Cat # 9212-0.25MG, Aldevron, Madison, WI) and Poly-L-Glutamic acid (PGA (15,000-50,000 kDa), 1.6 uL of 10 mg/mL solution in DNase free water, Cat # 26247-79-0, Millipore Sigma, Burlington, MA) for 15 minutes at 37*°*C to form the RNP complex. During incubation, T-cells were centrifuged for 300 g x 5 min and counted on the Countess II FL Automated Cell Counter (Thermo Fisher Scientific, Waltham, MA) with 0.4% Trypan Blue viability stain. One million cells per reaction were then aliquoted and spun for 90 g x 10 min. Following RNP incubation, linearized dsDNA HDR templates were added to the mixtures (2 uL) in PCR tubes and incubated for at least 5 minutes. Cells were then resuspended in 17.6 uL of P3 buffer (Lonza) and transferred to PCR tubes containing the RNP:DNA mixtures. Contents were then transferred to cuvettes (total volume 24 uL) and electroporated. Immediately following EP, 80 uL of Immunocult XF (STEMCELL Technologies, Vancouver, Canada) or TexMACS medium (Miltenyi Biotec, Bergisch Gladbach, Germany) with *no cytokines* was added to each reaction which were then rested for 30 minutes at 37*°*C. Cells were then moved to a flat-bottom 96 well plate containing 160 uL of medium supplemented with 500 U/mL IL-2 (Peprotech, Cranbury, NJ) or 10 ng/mL of IL-7 (BioTechne, Minneapolis, MN) and 10 ng/mL of IL-15 (BioTechne, Minneapolis, MN). Cells were cultured for 24 hours and then transferred to 12 well plates with 1 mL of media and incubated at 37 C for 48 hours. The TRAC guide sequence can be found in Supplementary Table 3.

### T-cell Culture

Metabolically-primed (MP) or Control *TRAC-*CAR T cells were cultured in TexMACS or Immunocult XF media supplemented with IL-7/IL-15 (10 ng/mL) or IL-2 (200 U/mL) at 1 million cells/mL respectively for the first three days. After EP, both MP and Control *TRAC-*CAR T cells were cultured in Immunocult XF for 7 days with IL-7/IL-15 (10 ng/mL) or IL-2 (500 U/mL). Every 2 days cells were centrifuged for 300 g x 5 min and counted on the Countess II FL Automated Cell Counter with 0.4% Trypan Blue viability stain. Cells were then resuspended in culture medium at 1 million cells/mL and the process repeated on days 5 and 7 post-EP.

### Scaled-Up GMP-compatible T-cell Manufacturing

For GMP-compatible experiments, T-cells were isolated and activated and cultured with either TransAct in TexMACS supplemented with IL-7 (10 ng/mL) or IL-15 (10 ng/mL) or with Immunocult XF activator and medium supplemented with 200 U/mL IL-2. Following activation, 20-50 million cells were electroporated on the CTS Xenon Electroporation System (ThermoFisher, Waltham, MA). TRAC sgRNA (1 uL/1E6 cells) (IDT, Coralville, IA) and SpCas9 (0.8 uL/1E6 cells) (Aldevron, Madison, WI) were mixed and incubated for 15 min at 37 C. Following incubation, linearized nanoplasmid template 1 uL/1E6 cells) was then added to the RNP mixtures. Cells were spun, counted, spun again for 5 minutes at 300 g, and resuspended in enough Genome Editing Buffer (Cat # A4998001, ThermoFisher, Waltham, MA) for a final volume of 1 mL when combined with the RNP:DNA mixture. Cells, DNA and RNPs were then added together, loaded into a Single-Shot cuvette (Cat # A50305, ThermoFisher, Waltham, MA), and electroporated on the Xenon unit at 1720 volts with a pulse width of 20 ms. Cells were then transferred to a T-25 flask with 4 mL of Immunocult XF medium containing no additives. After 30 minutes of rest at 37 C, Immunocult XF medium containing either IL-7 (10 ng/mL) with IL-15 (10 ng/mL) or 500 U/mL IL-2 was added to the T-75’s to bring the final concentration to 4 million cells/mL. After 24 hours, cells were transferred to a G-Rex 6M plate (10e6 per well) (Cat # P/N 80660M, Wilson Wolf, New Brighton, MN) and the final volume brought to 100 mL per well. Cells were then cultured at 37 C for 6 days after which 75 mL of media were aspirated, cells were collected, spun for 300 g x 5 min, and counted for use in endpoint assays.

### Flow Cytometry Analysis

CAR was detected using a 1A7 anti-14G2A antibody (National Cancer Institute, Biological Resources Branch) conjugated to APC using a Lightning Link APC Antibody Labeling kit (Cat # 705-0010, Novus Biologicals). TCR was detected using an anti-human TCR α/β antibody conjugate to BV421 (Biolegend, San Diego, CA). Flow cytometry to assess CAR and TCR positivity was performed on Day 8 of manufacturing on an Attune NxT flow cytometer (ThermoFisher, Waltham, MA). Immunophenotyping of cells was performed on Day 10 of manufacturing using a spectral immunophenotyping panel on an Aurora spectral cytometer (Cytek, California). Briefly, cells were plated in a 96-round bottom well plate (100k for CAR/TCR and 250k for spectral immunophenotyping), washed with 200 uL of PBS and spun at 1200 g x 1 min, twice. Cells were then stained for viability with either GhostRed 780 (Catalog # 50-105-2988, Tonbo Biosciences, San Diego, CA) or Live-Dead Blue (Catalog # L23105, ThermoFisher, Waltham MA). For CAR/TCR staining, 1 uL of Ghostred 780 was added to 10 mL of PBS to make a stock solution, 100 uL of stock solution was added to each sample and incubated for 30 minutes in the dark. For spectral flow staining, Live-Dead Blue stain was resuspended in 50 uL of DMSO, 1 uL added per 1 mL PBS to make a stock solution, and 200 uL of stock solution was added to each sample and incubated for 30 minutes in the dark. After viability stained, samples were washed twice and blocked for 30 minutes with 50 uL FACS buffer (0.5% BSA in PBS) with TruStain FcX solution (0.5 uL/sample) (Cat # 422301, Biolegend, San Diego, CA). Antibodies were then added to 100 uL of BD Brilliant Stain Buffer (Cat # 659611, BD Biosciences, Franklin Lakes, NJ) at the optimized amounts found in Supplementary Table 4 and incubated for 1 hour. Cells were then washed, resuspended in 200 or 75 uL of FACs buffer, and analyzed on the Attune or Aurora respectively. For spectral immunophenotyping, we used CD4, CD8, TCR, and CAR positivity to define populations and for all markers cells were gated by relative size, shape, singlets, viability, TCR negativity and CAR transgene positivity to find an analyzable population of viable CAR T cells. All antibodies are listed in Supplementary Table 4.

### *In Vitro* Cytotoxicity Assay on Incucyte

5,000 AkaLUC-GFP CHLA-20 cells were seeded in triplicate on 96-well plates and incubated for 24 hours at 37 C. 24 hours later 50,000, 25,000, or 10,000 CAR^+^ T-cells from Day 10 of manufacturing were added to each well for effector:target ratios of 5:1, 2.5:1, or 1:1. The plate was centrifuged for 5 minutes at 100g and then placed in the IncuCyte S3 Live-Cell Analysis System (Sartorius, G_ttingen, Germany) and stored at 37*°*C, 5% CO_2_. Images were taken every 3 hours for 48 hours. Green object count was used to calculate the number of cancer cells in each well and fluorescent images were analyzed with IncuCyte Base Analysis Software.

### *In vivo* Infusion into Mice with Human Xenografts

All animal experiments were approved by the University of Wisconsin-Madison Animal Care and Use Committee (ACUC protocol M005915). Male and female NOD-SCID-γc^-/-^ (NSG^TM^) mice (9-25 weeks old; Jackson Laboratory, Bar Harbor, ME) were subcutaneously injected with 10 million AkaLUC-GFP CHLA-20 GD2^+^ human neuroblastoma cells in the flank to establish tumors. After six days, tumor size was verified using bioluminescence measurements on the PerkinElmer *In Vivo* Imaging System (IVIS), and 3 million CAR^+^ T cells from Day 10 of manufacturing were injected into the tail vein of each mouse. Mice were imaged on the IVIS every 3-4 days after being sedated with Isoflurane and intraperitoneal injections of ∼120 mg/kg D-luciferin (GoldBio, St. Louis, MI). Mice were injected with 100,000 IU of human IL-2 (National Cancer Institute, Biological Resources Branch) subcutaneously on day 0 and following imaging. To quantify the total flux in images, a region of interest was drawn around established tumors on day 0 and calculated by Living Image Software (PerkinElmer; Total flux = radiance (photons/sec) in each pixel integrated over ROI area (cm^2^) x 4π). The minimum flux value was subtracted from each image to normalize for background signal.

### Flow cytometry of splenocytes

Spleens were removed, mechanically dissociated, and filtered using a Corning 70 uM cell strainer. Suspensions were centrifuged for 10 minutes at 300g and digested with ACK lysing buffer (Lonza, Walkerville, VA). The cells were then washed with PBS, centrifuged for 10 minutes at 300g, and resuspended in 1 mL of PBS. Cells were counted using Trypan Blue exclusion on the Countess II FL Automated Cell Counter. 1×10^6^ total cells were then added to 96-round well plates and stained for a spectral immunophenotyping panel for analysis on an Aurora spectral cytometer (Cytek). Briefly, samples were washed with PBS, stained with Live-Dead Blue for 30 minutes, blocked with FACS buffer and Trustain FcX, and incubated with the spectral immunophenotyping panel overnight. For spectral immunophenotyping, we used CD4, CD8, TCR, and CAR positivity to define populations and for all markers cells were gated by relative size, shape, singlets, viability, CD5 positivity, CD45 positivity, TCR negativity and CAR transgene positivity to find an analyzable population of viable CAR T cells. All antibodies and amounts are listed in Supplementary Table 4.

*In vitro* Serial Stimulation and Flow Cytometry Analysis of Lymphocytes: We added 120,000 CHLA-20 cells in duplicate to six well plates. We then seeded 120,000 CAR^+^ T-cells from Day 10 of manufacturing and incubated cells at 37*°*C, 5% CO_2_. 240,000 CHLA-20 cells were then added every 2-3 days for up to 20 days when the wells were harvested and stained for flow spectral cytometry. Briefly, CHLA-20/CAR T samples were washed with PBS, stained with Live-Dead Blue for 60 minutes, blocked with FACS buffer and Trustain FcX, and incubated with the spectral immunophenotyping panel overnight. For spectral immunophenotyping, we used TCR negativity to define populations and all markers cells were gated by relative size, shape, singlets, viability, CD5 positivity, CD45 positivity, and TCR negativity to find an analyzable population of viable T cells. All antibodies and amounts are listed in Supplementary Table 4.

### Cytokine Analysis

The supernatant of CAR T and cancer co-culture systems was measured for their expression of cytokines using the LegendPlex Human Essential Immune Response Panel (Cat # 740930, BioLegend, San Diego, CA) (IL-4, IL-2, CXCL10 (IP-10), IL-1β, TNF-α, CCL2 (MCP-1), IL-17A, IL-6, IL-10, IFN-γ, IL-12p70, CXCL8 (IL-8), TGF-β1 (Free Active Form). In brief, 50,000 AkaLUC-GFP CHLA-20 neuroblastoma cells were plated in 24 well plates with the addition of 250,000 CAR T cells 24 hours later. Suspension cells were harvested 24 hours later, spun for 5 minutes at 300g, cells counted, the supernatant flash frozen in liquid nitrogen, and stored at −20 C. Media samples were thawed and the manufacturer’s protocol for bead staining and analysis on an Attune NxT Cytometer (ThermoFisher, Waltham, MA) was followed. Briefly, standards for cytokines were prepared using 1:4 serial dilutions and co-culture supernatant added directly to a 96-well V-bottom plate. Mixed beads wereadded to each well and incubated on a plate mixer for 2 hours. The plate was then washed twice, detection antibody added to each well, incubated for 1 hour, and SA-PE was added to each well. The plate was incubated for 30 minutes, washed twice, and analyzed on the Attune Cytometer. Data was exported to LEGENDplex™ Data Analysis Software Suite where cytokine concentration was calculated. The data was exported to Excel, and cytokine concentration was normalized to the total number of T-cells in co-culture.

### Mitochondrial Staining

The mitochondrial mass and membrane potential of T-cells was measured by performing flow cytometry on cells stained with MitoTracker Green (Cat # M7514, ThermoFisher, Waltham, MA) and tetramethylrhodamine methyl ester perchlorate (TMRE) (ThermoFisher, Waltham, MA) dyes. Briefly, a stock solution of PBS containing 0.08 uL/mL of Mitotracker Green and 0.1 uL/mL of a 10 uM TMRE dye stock was created (200 uL per sample of 250,000 T-cells). Cells were stained for 20 minutes at 37*°*C, washed, and resuspended in 75 uL of FACS buffer (0.5% BSA in PBS).

### Metabolite Analysis

Media samples were taken from CAR T cell cultures on Day 10 of manufacturing and frozen at −20 C for future analysis. The Glucose-Glo and Lactate-Glo kits (Cat #’s J6021 and J5021, Promega, Madison, WI) were used to measure the apparent glucose and lactate concentrations in media samples according to the manufacturer’s protocol. Raw luminescence data were converted to concentration using the metabolite standards.

### Extracellular Flux Assay (Seahorse Assay)

The oxygen consumption rate (OCR) and extracellular acidification rate (ECAR) were measured following the manufacturer’s instructions for the Seahorse XF Cell Mito Stress Test Kit (Agilent, Madison, WI). Briefly, 5×10^5^ T-cells were resuspended in RPMI XF medium supplemented with 10mM glucose and 2mM glutamine and plated in a poly-L-lysine-coated XF96 plate. The T cell culture plate was centrifuged at 200g for 1 minute (no brake) and checked under the microscope to ensure even adhesion of T cells. T cells were then kept in a non-CO2 incubator for at least 1 hour before running the assay. The OCR and ECAR under basal conditions and in response to oligomycin (2.5 μM), fluorocarbonylcyanide-phenylhydrazone (FCCP) (1 μM), and rotenone/antimycin A (R/A) (0.5 μM) were measured using an XF96 Extracellular Flux Analyzer (Seahorse Bioscience, Madison, WI). Data was exported into Excel (V2311) using Agilent’s software and converted into graphs.

### Spectral Flow Cytometry Data Analysis

Analysis of spectral flow cytometry data was performed using Cytek’s SpectroFlo program., Single positive controls for each color were collected and analyzed in SpectroFlo for positive and negative populations. SpectroFlo’s unmixing algorithm was then used to compensate for spillover and autofluorescence of cells. Data was then exported to FlowJo (V10.9.0) where samples were gated for non-debris, singlets, and live cells. TCR, and CAR positivity were used to gate cell populations for *in vitro* samples and CD45, CD5, TCR, and CAR positivity for *in vivo* T-cell samples. Median fluorescent intensity (MFI)’s for each sample were calculated and input into Excel. Representative plots were generated in FlowJo using fluorescence minus one controls to set positive gates.

### Optical Metabolic Imaging

As previously described ^62, 63^, 200,000 CAR T cells in 75*μ*L fresh media were plated on Poly-D-Lysine coated 35mm glass bottom imaging dish (MatTek, Ashland,MA). Autofluorescence signals from NAD(P)H and FAD were imaged on a two-photon microscope (Ti-E, Nikon, Tokyo, Japan) at 750nm excitation (440/80nm emission) and 890nm excitation (550/50 emission), respectively. Fluorescence decay was collected with time-correlated single-photon counting electronics (SPC 150, Becker & Hickl GmbH) and fitted to a two-component exponential decay in SPCImage (v8.0, Becker and Hickl GmbH) to extract mean fluorescence lifetime (*τ*_m_) and fractional contribution of short- and long-lifetime components (*α*_1_ and *α*_2_). Single-cell nucleus and cytoplasm were manually segmented in CellProfiler and applied to the corresponding OMI images to quantify mean values of OMI parameters for each cell cytoplasm.

### Data Analysis and Software

All data analyses were performed in GraphPad Prism (V.10.0.2) and Microsoft Excel. Statistical tests were done in GraphPad Prism and indicated in the figure legends. Nanoplasmid sequences were designed in Benchling. FlowJo was used to analyze.fcs files exported from SpectroFlo and Attune NxT software. Representative flow plots were exported from FlowJo. UMAPs were created using the DownSample and UMAP_R plugins from to FlowJo to select for and cluster pooled data in a concatenated .fcs file. Figures were created and organized using Adobe Illustrator (V28.0). A p value less than 0.05 was defined as significant. Cohen’s effect size was calculated by dividing the difference between treated and control groups means and dividing by the pooled standard deviation of them.

## DATA AVAILBILITY STATEMENT

Data are available upon request.

## ACKNOWLEDGEMENTS

We thank members of the Saha, Capitini, Skala, Ayuso, Sodji labs, Kat Mueller, Tony Lauer, and Jolanta Vidugiriene for helpful discussion and comments on the manuscript, the University of Wisconsin (UW) Carbone Cancer Center Small Animal Imaging and Radiotherapy facility and Flow Cytometry Laboratory (supported by NIH P30 CA014520 and NIH S10 OD025225), Aldevron for technical support with Cas9 proteins, nanoplasmid preparation, and advice on linearization, Sarah Caroline Gomes de Lima for assistance in optimization experiments and helpful discussion, Amani Gillette for support and advice on Seahorse Experiments and mitochondrial staining, Paul Sondel (UW-Madison) and the National Cancer Institute for 1A7 anti-14G2a antibody for detection of CAR expression, James Thomson and Jue Zhang (Morgridge Institute for Research) for the AkaLUC-GFP CHLA20 cancer line used for in vivo studies, Promega for support on media sample analysis, ScaleReady for technical support on the G-Rex system, and ThermoFisher for support on optimization of EP on the CTS Xenon system.

We also acknowledge funding from the NIH R01 CA278051, the NSF Engineering Research Center (ERC) for Cell Manufacturing Technologies (CMaT) NSF-EEC 1648035 (MCS, CMC and KS), MACC Fund (CMC), St. Baldrick’s Foundation Empowering Pediatric Immunotherapy for Childhood Cancers Team grant, the UW-Madison Office of the Vice Chancellor for Research and Graduate Education with funding from the Wisconsin Alumni Research Foundation, Hyundai Hope on Wheels, the Grainger Institute for Engineering at UW-Madison (CMC and KS), and NIH R35 GM119644-01 (KS). The contents of this article do not necessarily reflect the views or policies of the Department of Health and Human Services, nor does mention of trade names, commercial products, or organizations imply endorsement by the US Government.

## AUTHOR CONTRIBUTIONS

These authors contributed equally: DC, DP, MHF. DC and MB developed the restriction digest and two-step purification procedure for HDR templates. DC, MB and AT optimized electroporation protocols. DC, MB, and AT isolated and cultured T cells and performed virus-free transfections. DP, BH, and LM assisted with cell culture. DC, MB, AT, BH, and LM performed flow cytometry. DC, DP and MHF performed cytokine experiments. MHF conducted in vivo experiments with support from QS. DC, DP, and MHF isolated mouse spleens/tumors and performed flow cytometry. DC, MB, and AT performed in vitro co-cultures. JV and TL performed analysis of metabolites in media. DC wrote the manuscript with input from all authors. DC, DP, MHF, MB, AT, JV, TL, BH, LM, and QS performed experiments and analyzed the data. DP provided integral initial insights that determined the direction of the study. KS, QS, MCS, and CMC supervised the research.

## REFERENCES

1. Chen, Y.-J., Abila, B., and Mostafa Kamel, Y. (2023). CAR-T: What is next? Cancers (Basel) 15, 663. 10.3390/cancers15030663.

2. Zhang, C., Liu, J., Zhong, J.F., and Zhang, X. (2017). Engineering CAR-T cells. Biomark Res 5, 22. 10.1186/s40364-017-0102-y.

3. June, C.H., O’Connor, R.S., Kawalekar, O.U., Ghassemi, S., and Milone, M.C. (2018). CAR T cell immunotherapy for human cancer. Science 359, 1361–1365. 10.1126/science.aar6711.

4. Louis, C.U., Savoldo, B., Dotti, G., Pule, M., Yvon, E., Myers, G.D., Rossig, C., Russell, H.V., Diouf, O., Liu, E., et al. (2011). Antitumor activity and long-term fate of chimeric antigen receptor-positive T cells in patients with neuroblastoma. Blood 118, 6050–6056. 10.1182/blood-2011-05-354449.

5. Pule, M.A., Savoldo, B., Myers, G.D., Rossig, C., Russell, H.V., Dotti, G., Huls, M.H., Liu, E., Gee, A.P., Mei, Z., et al. (2008). Virus-specific T cells engineered to coexpress tumor-specific receptors: persistence and antitumor activity in individuals with neuroblastoma. Nat. Med. 14, 1264–1270. 10.1038/nm.1882.

6. Del Bufalo, F., De Angelis, B., Caruana, I., Del Baldo, G., De Ioris, M.A., Serra, A., Mastronuzzi, A., Cefalo, M.G., Pagliara, D., Amicucci, M., et al. (2023). GD2-CART01 for Relapsed or Refractory High-Risk Neuroblastoma. N. Engl. J. Med. 388, 1284–1295. 10.1056/NEJMoa2210859.

7. Reddy, O.L., Stroncek, D.F., and Panch, S.R. (2020). Improving CAR T cell therapy by optimizing critical quality attributes. Semin. Hematol. 57, 33–38. 10.1053/j.seminhematol.2020.07.005.

8. Tantalo, D.G., Oliver, A.J., von Scheidt, B., Harrison, A.J., Mueller, S.N., Kershaw, M.H., and Slaney, C.Y. (2021). Understanding T cell phenotype for the design of effective chimeric antigen receptor T cell therapies. J Immunother Cancer 9. 10.1136/jitc-2021-002555.

9. Restifo, N.P., and Gattinoni, L. (2013). Lineage relationship of effector and memory T cells. Curr. Opin. Immunol. 25, 556–563. 10.1016/j.coi.2013.09.003.

10. Piscopo, N.J., Mueller, K.P., Das, A., Hematti, P., Murphy, W.L., Palecek, S.P., Capitini, C.M., and Saha, K. (2018). Bioengineering Solutions for Manufacturing Challenges in CAR T Cells. Biotechnol. J. 13. 10.1002/biot.201700095.

11. Yu, J., Chen, P., Watson, A., Truong, D., Antonchuk, J., Kokaji, A.I., Woodside, S.M., Eaves, A.C., and Thomas, T.E. (2019). Optimization of human T cell activation and expansion protocols improves ef ciency of genetic Modi cation and overall cell yield. https://cdn.stemcell.com/media/files/poster/SP00235-Optimization_of_Human_T_Cell_Activation_and_Expansion_Protocols_Improves_Efficiency_of_Genetic_Modification_and_Overall_Cell_Yield.pdf.

12. Wan, Y.Y., and Flavell, R.A. (2009). How Diverse—CD4 Effector T Cells and their Functions. J. Mol. Cell Biol. 1, 20–36. 10.1093/jmcb/mjp001.

13. Gattinoni, L., Lugli, E., Ji, Y., Pos, Z., Paulos, C.M., Quigley, M.F., Almeida, J.R., Gostick, E., Yu, Z., Carpenito, C., et al. (2011). A human memory T cell subset with stem cell–like properties. Nat. Med. 17, 1290–1297. 10.1038/nm.2446.

14. Biasco, L., Izotova, N., Rivat, C., Ghorashian, S., Richardson, R., Guvenel, A., Hough, R., Wynn, R., Popova, B., Lopes, A., et al. (2021). Clonal expansion of T memory stem cells determines early anti-leukemic responses and long-term CAR T cell persistence in patients. Nat Cancer 2, 629–642. 10.1038/s43018-021-00207-7.

15. Ghassemi, S., Prachi, P., Scholler, J., Nunez-Cruz, S., Barrett, D.M., Bedoya, F., Fraietta, J.A., Lacey, S.F., Levine, B.L., Grupp, S.A., et al. (2016). Minimally Ex Vivo Manipulated Gene-Modified T Cells Display Enhanced Tumor Control. Blood 128, 4549–4549. 10.1182/blood.V128.22.4549.4549.

16. Ghassemi, S., Martinez-Becerra, F.J., Master, A.M., Richman, S.A., Heo, D., Leferovich, J., Tu, Y., García-Cañaveras, J.C., Ayari, A., Lu, Y., et al. (2020). Enhancing Chimeric Antigen Receptor T Cell Anti-tumor Function through Advanced Media Design. Mol Ther Methods Clin Dev 18, 595– 606. 10.1016/j.omtm.2020.07.008.

17. MacPherson, S., Keyes, S., Kilgour, M.K., Smazynski, J., Chan, V., Sudderth, J., Turcotte, T., Devlieger, A., Yu, J., Huggler, K.S., et al. (2022). Clinically relevant T cell expansion media activate distinct metabolic programs uncoupled from cellular function. Molecular Therapy - Methods & Clinical Development 24, 380–393. 10.1016/j.omtm.2022.02.004.

18. Raud, B., McGuire, P.J., Jones, R.G., Sparwasser, T., and Berod, L. (2018). Fatty acid metabolism in CD8+ T cell memory: Challenging current concepts. Immunol. Rev. 283, 213–231. 10.1111/imr.12655.

19. Chang, C.-H., Curtis, J.D., Maggi, L.B., Jr, Faubert, B., Villarino, A.V., O’Sullivan, D., Huang, S.C.-C., van der Windt, G.J.W., Blagih, J., Qiu, J., et al. (2013). Posttranscriptional control of T cell effector function by aerobic glycolysis. Cell 153, 1239–1251. 10.1016/j.cell.2013.05.016.

20. Pearce, E.L., Walsh, M.C., Cejas, P.J., Harms, G.M., Shen, H., Wang, L.-S., Jones, R.G., and Choi, Y. (2009). Enhancing CD8 T-cell memory by modulating fatty acid metabolism. Nature 460, 103–107. 10.1038/nature08097.

21. Zhou, J., Jin, L., Wang, F., Zhang, Y., Liu, B., and Zhao, T. (2019). Chimeric antigen receptor T (CAR-T) cells expanded with IL-7/IL-15 mediate superior antitumor effects. Protein Cell 10, 764– 769. 10.1007/s13238-019-0643-y.

22. van der Windt, G.J.W., and Pearce, E.L. (2012). Metabolic switching and fuel choice during T-cell differentiation and memory development. Immunol. Rev. 249, 27–42. 10.1111/j.1600-065X.2012.01150.x.

23. Le Bourgeois, T., Strauss, L., Aksoylar, H.-I., Daneshmandi, S., Seth, P., Patsoukis, N., and Boussiotis, V.A. (2018). Targeting T Cell Metabolism for Improvement of Cancer Immunotherapy. Front. Oncol. 8, 237. 10.3389/fonc.2018.00237.

24. Dugnani, E., Pasquale, V., Bordignon, C., Canu, A., Piemonti, L., and Monti, P. (2017). Integrating T cell metabolism in cancer immunotherapy. Cancer Lett. 411, 12–18. 10.1016/j.canlet.2017.09.039.

25. Shen, L., Xiao, Y., Zhang, C., Li, S., Teng, X., Cui, L., Liu, T., Wu, N., and Lu, Z. (2022). Metabolic reprogramming by ex vivo glutamine inhibition endows CAR-T cells with less-differentiated phenotype and persistent antitumor activity. Cancer Lett. 538, 215710. 10.1016/j.canlet.2022.215710.

26. Nabe, S., Yamada, T., Suzuki, J., Toriyama, K., Yasuoka, T., Kuwahara, M., Shiraishi, A., Takenaka, K., Yasukawa, M., and Yamashita, M. (2018). Reinforce the antitumor activity of CD8+ T cells via glutamine restriction. Cancer Sci. 109, 3737–3750. 10.1111/cas.13827.

27. Klein Geltink, R.I., Edwards-Hicks, J., O’Sullivan, D., Sanin, D.E., Patterson, A.E., Puleston, D.J., Ligthart, N.A.M., Buescher, J.M., Grzes, K.M., Kabat, A.M., et al. (2020). Metabolic conditioning of CD8+ effector T cells for adoptive cell therapy. Nature Metabolism 2, 703–716. 10.1038/s42255-020-0256-z.

28. Amini, A., and Veraitch, F. (2019). Glucose deprivation enriches for central memory T cells during chimeric antigen receptor-T cell expansion. Cytotherapy 21, S30–S31. 10.1016/j.jcyt.2019.03.348.

29. Pilipow, K., Scamardella, E., Puccio, S., Gautam, S., De Paoli, F., Mazza, E.M., De Simone, G., Polletti, S., Buccilli, M., Zanon, V., et al. (2018). Antioxidant metabolism regulates CD8+ T memory stem cell formation and antitumor immunity. JCI Insight 3. 10.1172/jci.insight.122299.

30. Cepko, C., and Pear, W. (2001). Overview of the retrovirus transduction system. Curr. Protoc. Mol. Biol. Chapter 9, Unit9.9. 10.1002/0471142727.mb0909s36.

31. Watanabe, N., and McKenna, M.K. (2022). Generation of CAR T-cells using γ-retroviral vector. Methods Cell Biol. 167, 171–183. 10.1016/bs.mcb.2021.06.014.

32. Ajina, A., and Maher, J. (2018). Strategies to Address Chimeric Antigen Receptor Tonic Signaling. Mol. Cancer Ther. 17, 1795–1815. 10.1158/1535-7163.MCT-17-1097.

33. Shams, F., Bayat, H., Mohammadian, O., Mahboudi, S., Vahidnezhad, H., Soosanabadi, M., and Rahimpour, A. (2022). Advance trends in targeting homology-directed repair for accurate gene editing: An inclusive review of small molecules and modified CRISPR-Cas9 systems. Bioimpacts 12, 371–391. 10.34172/bi.2022.23871.

34. Fu, Y.-W., Dai, X.-Y., Wang, W.-T., Yang, Z.-X., Zhao, J.-J., Zhang, J.-P., Wen, W., Zhang, F., Oberg, K.C., Zhang, L., et al. (2021). Dynamics and competition of CRISPR-Cas9 ribonucleoproteins and AAV donor-mediated NHEJ, MMEJ and HDR editing. Nucleic Acids Res. 49, 969–985. 10.1093/nar/gkaa1251.

35. Webber, B.R., Johnson, M.J., Skeate, J.G., Slipek, N.J., Lahr, W.S., DeFeo, A.P., Mills, L.J., Qiu, X., Rathmann, B., Diers, M.D., et al. (2023). Cas9-induced targeted integration of large DNA payloads in primary human T cells via homology-mediated end-joining DNA repair. Nat Biomed Eng. 10.1038/s41551-023-01157-4.

36. Foy, S.P., Jacoby, K., Bota, D.A., Hunter, T., Pan, Z., Stawiski, E., Ma, Y., Lu, W., Peng, S., Wang, C.L., et al. (2022). Non-viral precision T cell receptor replacement for personalized cell therapy. Nature, 1–10. 10.1038/s41586-022-05531-1.

37. Hu, Y., Zu, C., Zhang, M., Wei, G., Li, W., Fu, S., Hong, R., Zhou, L., Wu, W., Cui, J., et al. (2023). Safety and efficacy of CRISPR-based non-viral PD1 locus specifically integrated anti-CD19 CAR-T cells in patients with relapsed or refractory Non-Hodgkin’s lymphoma: a first-in-human phase I study. EClinicalMedicine 60, 102010. 10.1016/j.eclinm.2023.102010.

38. Ye, L., Park, J.J., Peng, L., Yang, Q., Chow, R.D., Dong, M.B., Lam, S.Z., Guo, J., Tang, E., Zhang, Y., et al. (2022). A genome-scale gain-of-function CRISPR screen in CD8 T cells identifies proline metabolism as a means to enhance CAR-T therapy. Cell Metab. 34, 595–614.e14. 10.1016/j.cmet.2022.02.009.

39. Eyquem, J., Mansilla-Soto, J., Giavridis, T., van der Stegen, S.J.C., Hamieh, M., Cunanan, K.M., Odak, A., Gönen, M., and Sadelain, M. (2017). Targeting a CAR to the TRAC locus with CRISPR/Cas9 enhances tumour rejection. Nature 543, 113–117. 10.1038/nature21405.

40. Stadtmauer, E.A., Fraietta, J.A., Davis, M.M., Cohen, A.D., Weber, K.L., Lancaster, E., Mangan, P.A., Kulikovskaya, I., Gupta, M., Chen, F., et al. (2020). CRISPR-engineered T cells in patients with refractory cancer. Science 367. 10.1126/science.aba7365.

41. Mueller, K.P., Piscopo, N.J., Forsberg, M.H., Saraspe, L.A., Das, A., Russell, B., Smerchansky, M., Cappabianca, D., Shi, L., Shankar, K., et al. (2022). Production and characterization of virus-free, CRISPR-CAR T cells capable of inducing solid tumor regression. J Immunother Cancer 10. 10.1136/jitc-2021-004446.

42. Pham, D.L., Cappabianca, D., Forsberg, M.H., Weaver, C., Mueller, K.P., Tommasi, A., Vidugiriene, J., Lauer, A., Sylvester, K., Bugel, M., et al. (2024). Label free metabolic imaging to enhance the efficacy of Chimeric Antigen Receptor T cell therapy. bioRxiv, 2024.02.20.581240. 10.1101/2024.02.20.581240.

43. Wang, X., and Rivière, I. (2016). Clinical manufacturing of CAR T cells: foundation of a promising therapy. Mol Ther Oncolytics 3, 16015. 10.1038/mto.2016.15.

44. Gattinoni, L., Speiser, D.E., Lichterfeld, M., and Bonini, C. (2017). T memory stem cells in health and disease. Nat. Med. 23, 18–27. 10.1038/nm.4241.

45. Xu, Y., Zhang, M., Ramos, C.A., Durett, A., Liu, E., Dakhova, O., Liu, H., Creighton, C.J., Gee, A.P., Heslop, H.E., et al. (2014). Closely related T-memory stem cells correlate with in vivo expansion of CAR.CD19-T cells and are preserved by IL-7 and IL-15. Blood 123, 3750–3759. 10.1182/blood-2014-01-552174.

46. Fritsch, R.D., Shen, X., Sims, G.P., Hathcock, K.S., Hodes, R.J., and Lipsky, P.E. (2005). Stepwise differentiation of CD4 memory T cells defined by expression of CCR7 and CD27. J. Immunol. 175, 6489–6497. 10.4049/jimmunol.175.10.6489.

47. Desdín-Micó, G., Soto-Heredero, G., and Mittelbrunn, M. (2018). Mitochondrial activity in T cells. Mitochondrion 41, 51–57. 10.1016/j.mito.2017.10.006.

48. Atkins, R.M., Menges, M.A., Bauer, A., Turner, J.G., and Locke, F.L. (2020). Metabolically Flexible CAR T Cells (mfCAR-T), with Constitutive Expression of PGC-1α Resistant to Post Translational Modifications, Exhibit Superior Survival and Function in Vitro. Blood 136, 30. 10.1182/blood-2020-143217.

49. Claiborne, M.D. (2022). Manipulation of metabolic pathways to promote stem-like and memory T cell phenotypes for immunotherapy. Front. Immunol. 13, 1061411. 10.3389/fimmu.2022.1061411.

50. Moore, T.V., Clay, B.S., Cannon, J.L., Histed, A., Shilling, R.A., and Sperling, A.I. (2011). Inducible costimulator controls migration of T cells to the lungs via down-regulation of CCR7 and CD62L. Am. J. Respir. Cell Mol. Biol. 45, 843–850. 10.1165/rcmb.2010-0466OC.

51 Portero-Sainz, I., Gómez-García de Soria, V., Cuesta-Mateos, C., Fernández-Arandojo, C., Vega-Piris, L., Royg, M., Colom-Fernández, B., Marcos-Jiménez, A., Somovilla-Crespo, B., Ramírez-Mengíbar, A., et al. (2017). A high migratory capacity of donor T-cells in response to the lymph node homing receptor CCR7 increases the incidence and severity of GvHD. Bone Marrow Transplant. 52, 745–752. 10.1038/bmt.2016.342.

52. Yamazaki, K. (2002). An ultrastructural study of cutaneous panniculitis-like T-cell lymphoma: cytoplasmic granules and active cellular and cell-to-matrix interaction mimic cytotoxic T-cells. Ultrastruct. Pathol. 26, 185–190. 10.1080/01913120290076856.

53. Golubovskaya, V., and Wu, L. (2016). Different Subsets of T Cells, Memory, Effector Functions, and CAR-T Immunotherapy. Cancers 8. 10.3390/cancers8030036.

54. Schwab, R.D., Bedoya, D.M., King, T.R., Levine, B.L., and Posey, A.D., Jr (2020). Approaches of T Cell Activation and Differentiation for CAR-T Cell Therapies. Methods Mol. Biol. 2086, 203–211. 10.1007/978-1-0716-0146-4_15.

55. Balke-Want, H., Keerthi, V., Cadinanos-Garai, A., Fowler, C., Gkitsas, N., Brown, A.K., Tunuguntla, R., Abou-El-Enein, M., and Feldman, S.A. (2023). Non-viral chimeric antigen receptor (CAR) T cells going viral. Immunooncol Technol 18, 100375. 10.1016/j.iotech.2023.100375.

56. Prather, K.J., Sagar, S., Murphy, J., and Chartrain, M. (2003). Industrial scale production of plasmid DNA for vaccine and gene therapy: plasmid design, production, and purification. Enzyme Microb. Technol. 33, 865–883. 10.1016/S0141-0229(03)00205-9.

57. Listner, K., Bentley, L., Okonkowski, J., Kistler, C., Wnek, R., Caparoni, A., Junker, B., Robinson, D., Salmon, P., and Chartrain, M. (2006). Development of a highly productive and scalable plasmid DNA production platform. Biotechnol. Prog. 22, 1335–1345. 10.1021/bp060046h.

58. Fitzgerald, K.C., Bhargava, P., Smith, M.D., Vizthum, D., Henry-Barron, B., Kornberg, M.D., Cassard, S.D., Kapogiannis, D., Sullivan, P., Baer, D.J., et al. (2022). Intermittent calorie restriction alters T cell subsets and metabolic markers in people with multiple sclerosis. EBioMedicine 82, 104124. 10.1016/j.ebiom.2022.104124.

59. Collins, N., Han, S.-J., Enamorado, M., Link, V.M., Huang, B., Moseman, E.A., Kishton, R.J., Shannon, J.P., Dixit, D., Schwab, S.R., et al. (2019). The Bone Marrow Protects and Optimizes Immunological Memory during Dietary Restriction. Cell 178, 1088–1101.e15. 10.1016/j.cell.2019.07.049.

60. Chen, Y., Xu, Z., Sun, H., Ouyang, X., Han, Y., Yu, H., Wu, N., Xie, Y., and Su, B. (2023). Regulation of CD8+ T memory and exhaustion by the mTOR signals. Cell. Mol. Immunol. 20, 1023–1039. 10.1038/s41423-023-01064-3.

61. Carnes, A., Tiwari, N., Beilowitz, J., Sampson, C., Peterson, D., and Williams, J. (2016). Production of a Nanoplasmid^TM^ with a large gene insert using the HyperGRO^TM^ fermentation process.

62. Walsh, A.J., Cook, R.S., Manning, H.C., Hicks, D.J., Lafontant, A., Arteaga, C.L., and Skala, M.C. (2013). Optical metabolic imaging identifies glycolytic levels, subtypes, and early-treatment response in breast cancer. Cancer Res. 73, 6164–6174. 10.1158/0008-5472.CAN-13-0527.

63. Walsh, A.J., Mueller, K., Jones, I., Heaster, T.M., Saha, K., and Skala, M.C. (2018). Autofluorescence Imaging of T cell Activation. The Journal of Immunology 200, 120–113.

