## Supplemental Materials and Tables for "Metabolic priming of GD2 *TRAC*-CAR T cells during manufacturing promotes memory phenotypes while enhancing persistence"

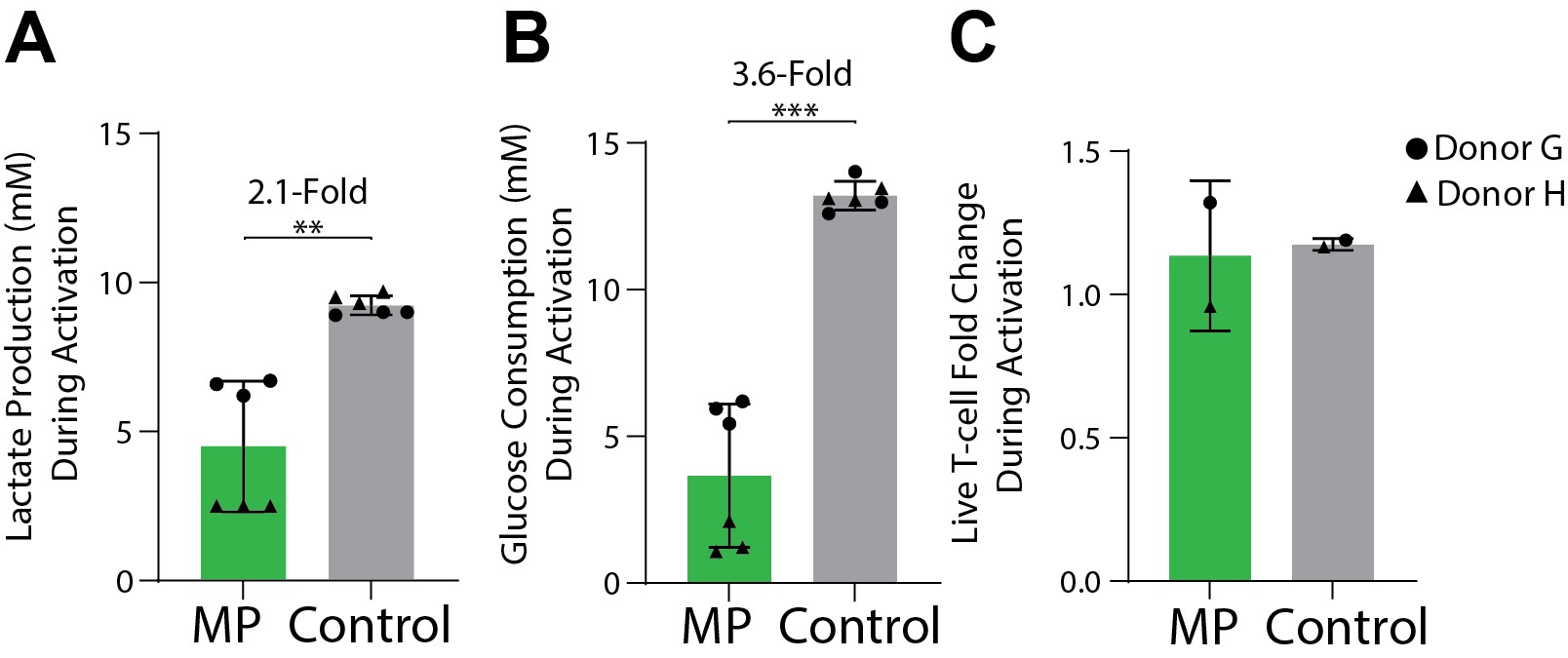

**Figure S1. Effect of Activation Conditions on Lactate Production and Glucose Consumption.**  (A) Lactate production, (B) glucose consumption and (C) proliferation of MP or Control T cells during activation (MP – 72 hours; Control – 48 hours). 2 donors, (lactate/glucose) N_MP_ = N_Control_ = 6, (proliferation) N_MP_ = N_Control_ = 2. Error bars represent mean and standard deviation. Statistical significance was determined with a paired-test; **p<0.01; ***p<0.001.

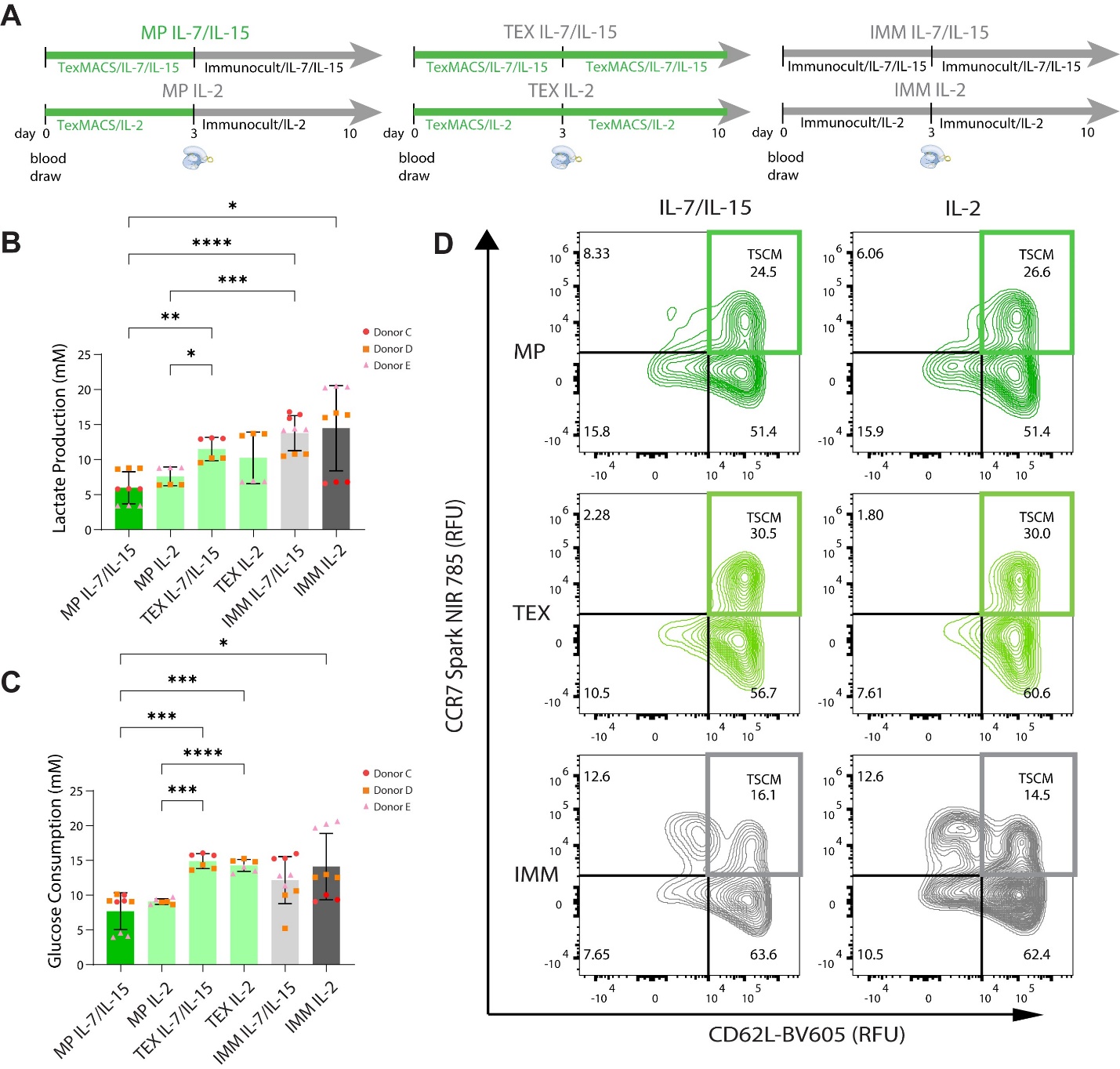

**Figure S2. Stem Cell Memory Phenotypes of Differentially Cultured *TRAC-*CAR T cells. (A)** CAR T cells were manufactured under six different media/cytokine conditions: 1). Metabolic priming with IL-7/IL-15 (MP), 2). metabolic priming with IL-2, 3). TexMACs with IL-7/IL-15, 4). TexMACs with IL-2, 5). Immunocult with IL-7/IL-15, and 6). Immunocult with IL-2 (Ctrl). Bar graphs of **(B)** lactate production (MP IL-7/I,-15: 6.0 mM (2.3), MP IL-2: 7.6 (1.3), TEX IL-7/IL-15: 11.5 (1.7), TEX IL-2: 10.3 (3.7), IMM IL-7/IL-15: 13.8 (2.5), IMM IL-2: 14.5 (6.1), *p* (IMM IL-2 vs MP IL-7/IL-15) = 0.0341, *p* (TEX IL-7/IL-15 vs MP IL-2) = 0.0147, *p* (TEX IL-7/IL-15 vs MP IL-7/IL-15) = 0.0017, *p* (IMM IL-7/IL-15 vs MP IL-2 or IL-7/IL-15) < 0.001. and **(C)** glucose consumption (MP IL-7/I,-15: 7.7 (2.6) , MP IL-2: 9.1 (0.4), TEX IL-7/IL-15: 14.9 (1.1), TEX IL-2: 14.3 (0.9), IMM IL-7/IL-15: 12.2 (3.4), IMM IL-2: 14.1 (4.8), *p* (IMM IL-2 vs MP IL-7/IL-15) = 0.046, *p* (TEX IL-2 or IL-7/IL-15 vs MP IL-2 or IL-7/IL-15) < 0.001 for all) from day 8 to 10 of manufacturing of differentially-cultured *TRAC-*CAR T cells. **(D)** Representative contour plots of the expression of CCR7 vs CD62L on Day 10 of manufacturing. 2-3 donors, N_MP, IL-7/IL-15_ = 9, N_MP, IL-2_ = 6, N_TEX, IL-7/IL-15_ = 6, N_TEX, IL-2_ = 6, N_IMM, IL-7/IL-15_ = 9, N_IMM, IL-2_ = 9. Error bars represent mean and standard deviation. Statistical significance was determined with Brown-Forsythe and Welch ANOVA tests using Dunnett’s T3 test for multiple comparisons; *p<0.05; **p<0.01; ***p<0.001; ****p<0.0001.

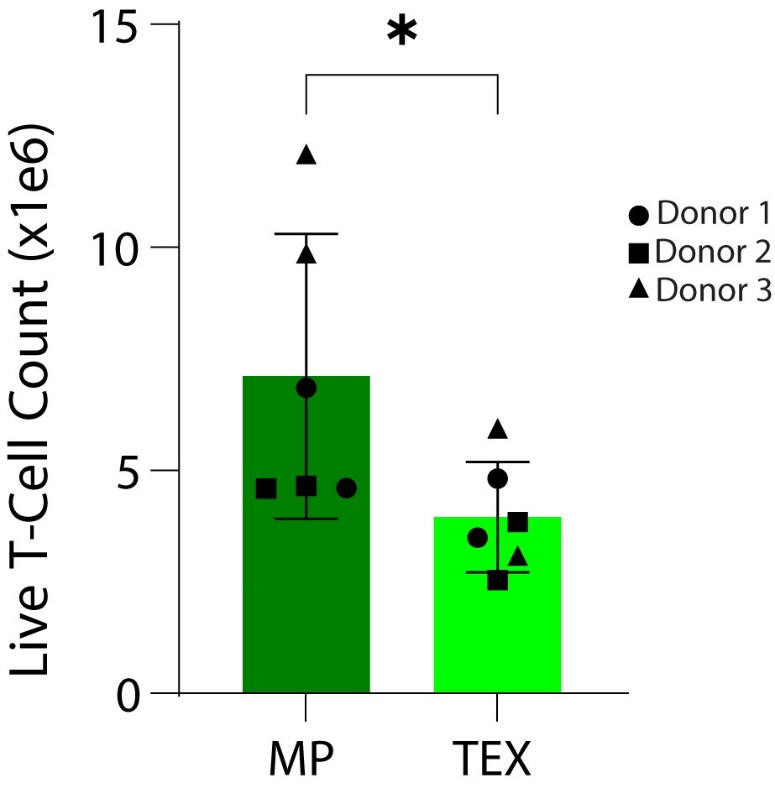

**Figure S3. *TRAC-*CAR T Cell Expansion Under MP and TEX Cultures Conditions.** *TRAC-*CAR T cells were manufactured with MP (metabolic priming) or TEX (TexMACs media only) culture, both supplemented with IL-7/IL-15. Bar graph of the live T-cell count on Day 5 post-EP of manufacturing (MP: 7.2e6 cells (3.2), TEX: 4.0e6 cells (1.2), *p* = 0.041). 3 donors, N_MP_ = 6, N_TEX_ = 6. Error bars represent mean and standard deviation. Statistical significance was determined with paired t-tests; *p<0.05.

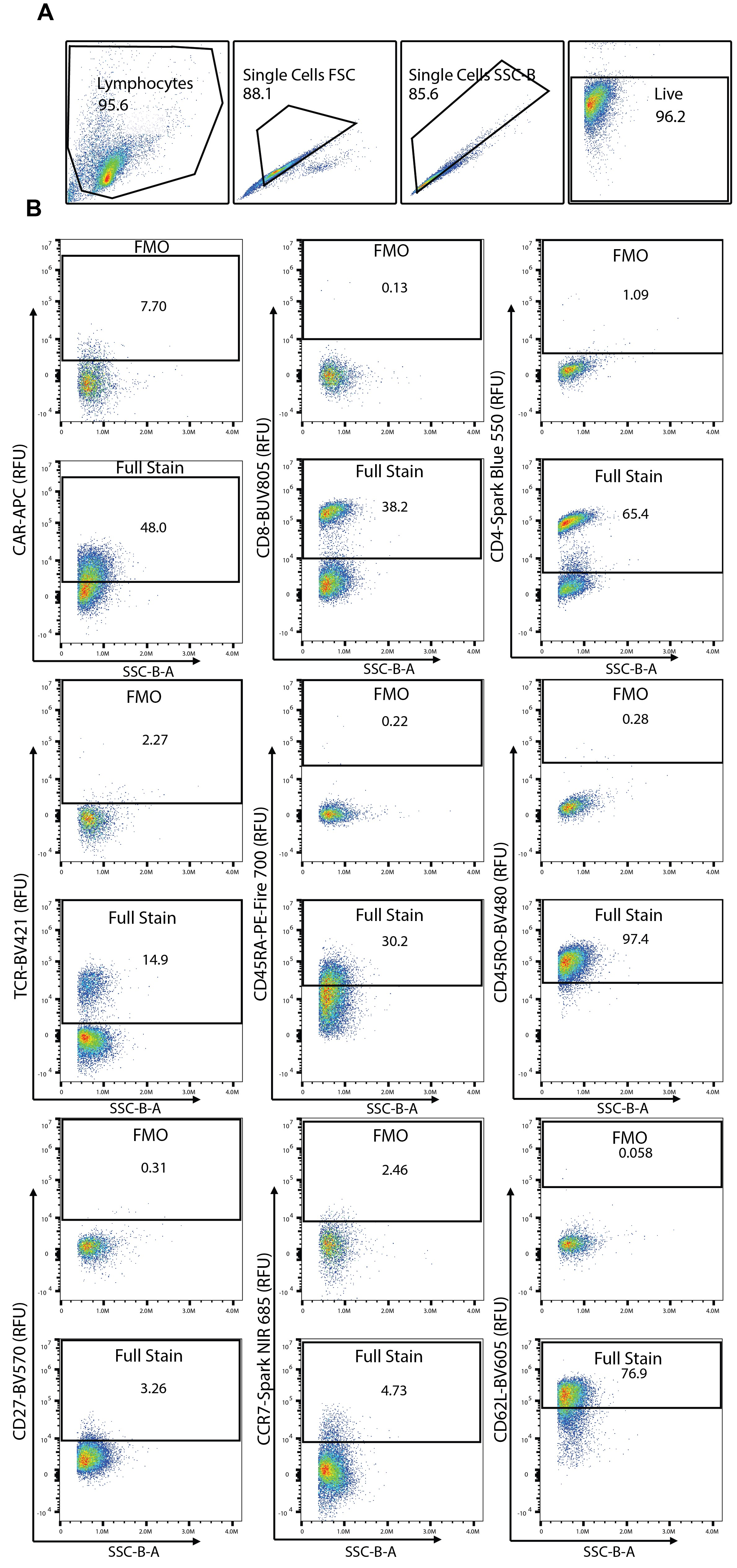

**Figure S4. Post-Manufacturing CAR T Cell Gating Strategy. (A)** Gating strategy for analysis of spectral immunophenotyping flow cytometry data of CAR T cells post-manufacturing. **(B)** FMO’s and representative positive populations depicting positive and negative gates.

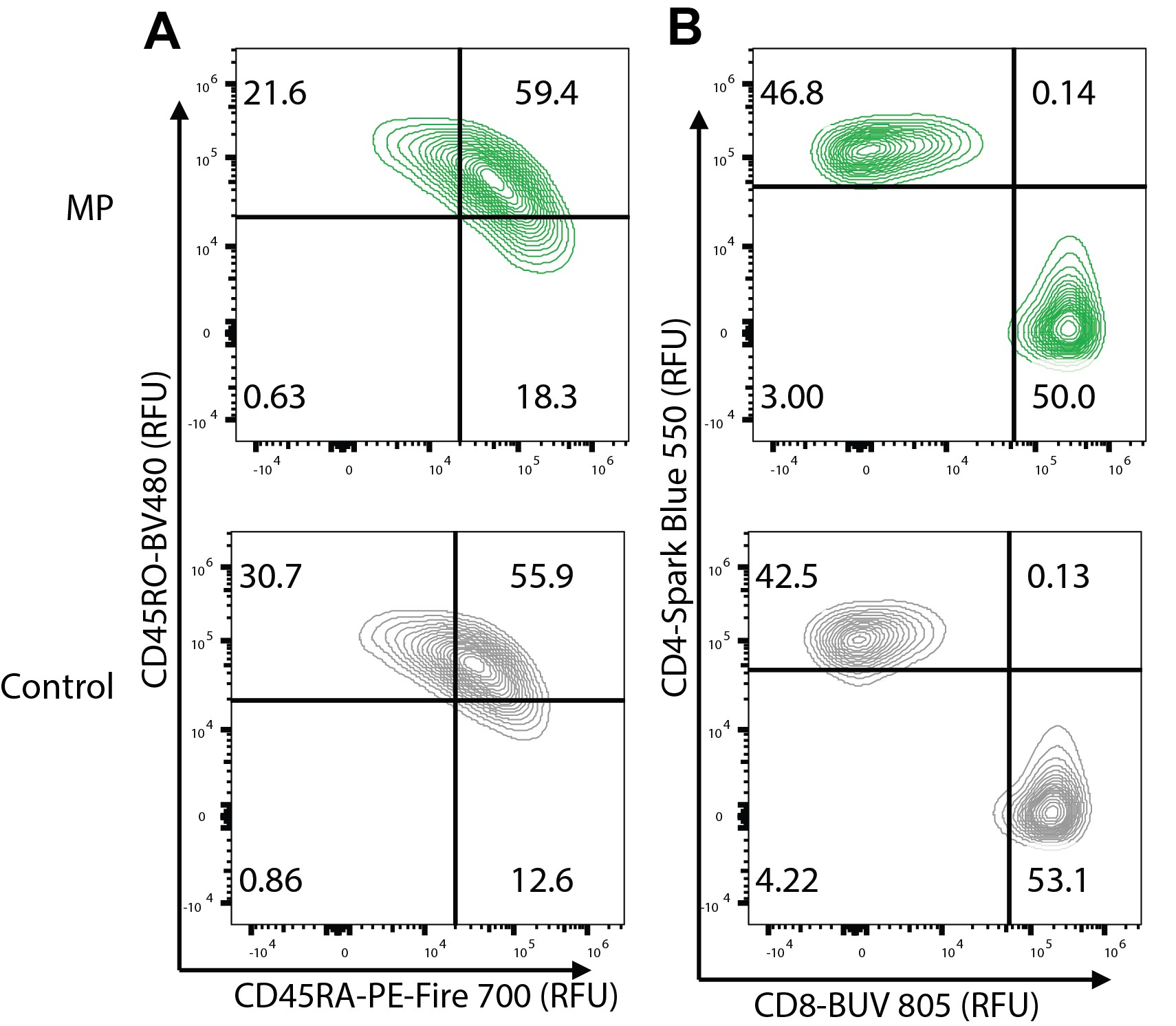

**Figure S5. Expression of CD45RA/CD45RO and CD4/CD8 in MP *TRAC-*CAR T Cells at Scale.** Representative contour plots for expression of (A) CD45RA/CD45RO and (B) CD4/CD8 MP or Control *TRAC-*CAR T cells (Donor F shown).

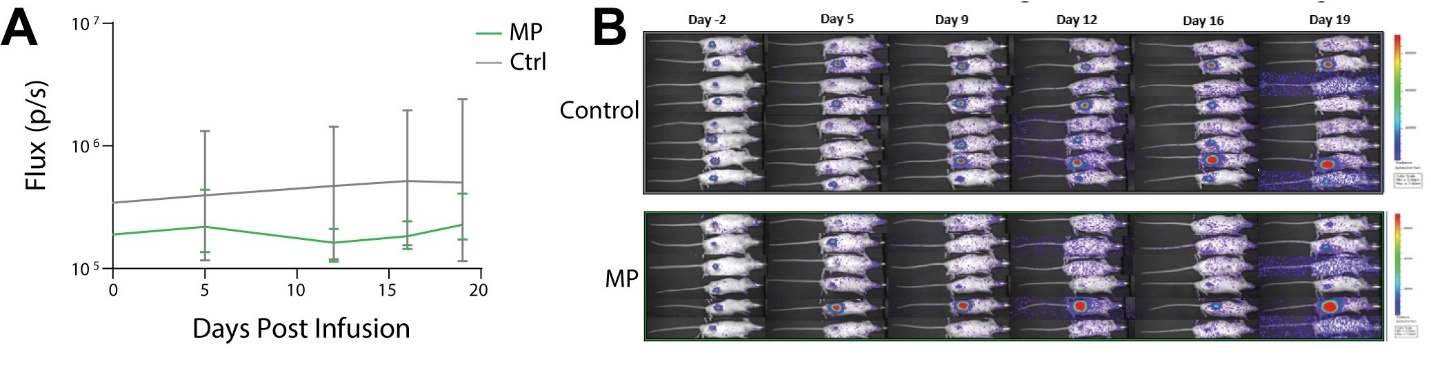

**Figure S6. In Vivo Potency of MP and Control TRAC-CAR T cells. (A)** The flux over time (luminescence from IVIS images) is shown for mice treated with MP or Control *TRAC*-CAR T cells. **(B)** IVIS images depict tumor growth over time for the same conditions. (2 donors, N_MP_ = 6, N_Control_ = 8). Error bars represent mean and standard deviation.

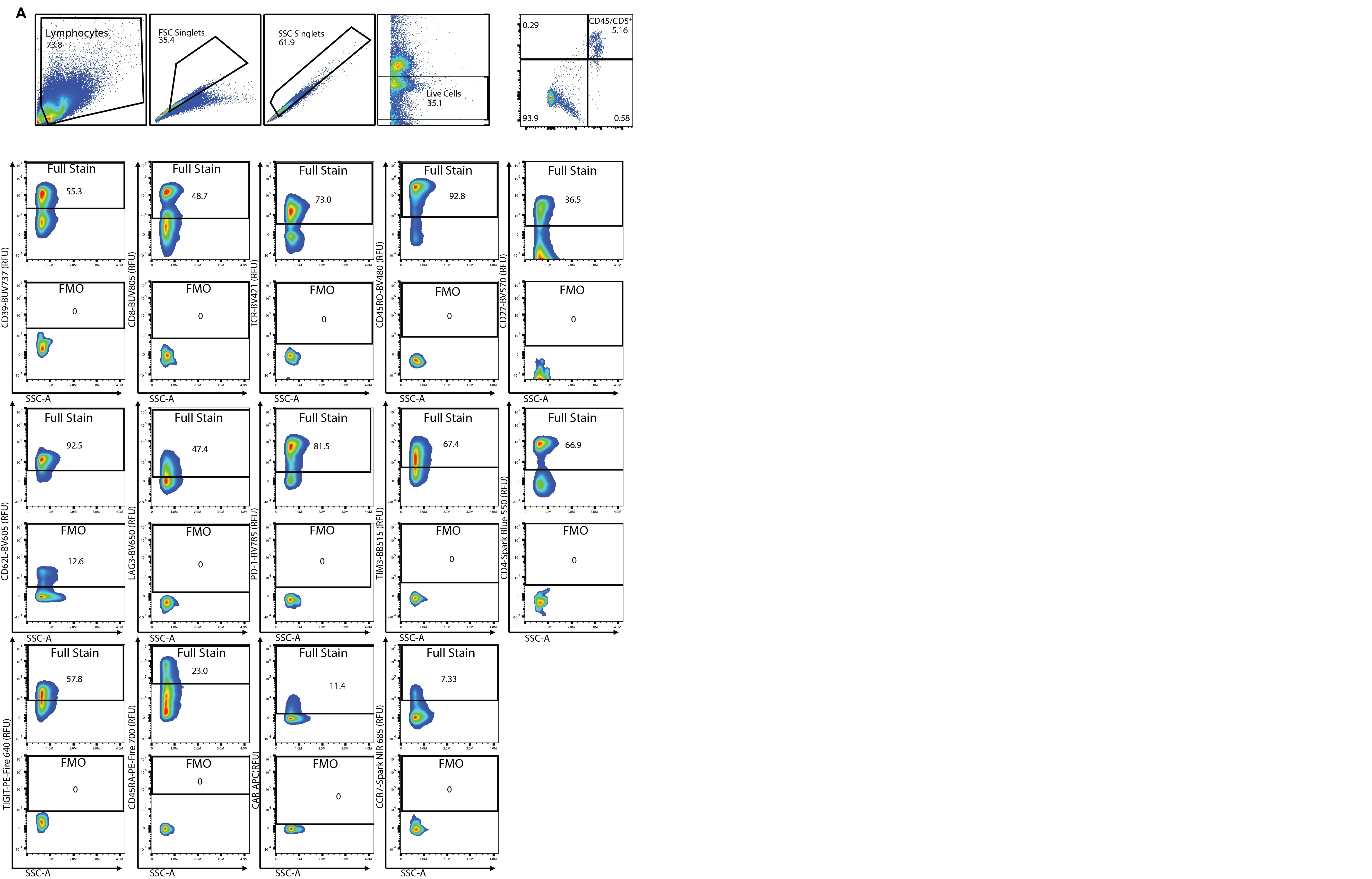

**Figure S7 Gating Scheme for Analysis of Lymphocytes in Mouse Spleens.** Gating strategy for analysis of live, CD5/CD45^+^ lymphocytes in isolated mouse spleens. **(B)** FMO’s and representative positive populations depicting positive and negative gates.

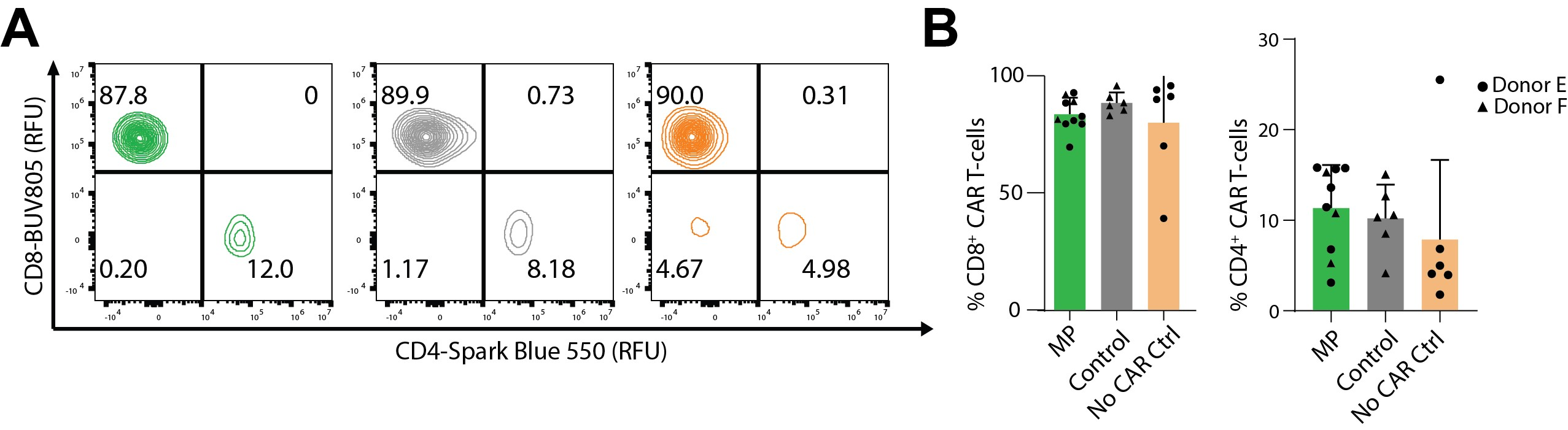
**Figure S8. Expression of CD4 and CD8 Post In Vivo Tumor Challenge. (A)** Representative contour plots for CD4/CD8 expression in CD5+/CD45+/TCR-/transgene+ lymphocytes isolated from mouse spleens (Green = MP, Donor E; Gray = Control, Donor F; Orange = No CAR Ctrl, Donor E). **(B)** Bar graphs for relative expression of CD4 and CD8 for MP or Control -CAR T cells and no CAR control T-cells. Samples with less than 20 CD5+/CD45+/TCR-/transgene+ events were excluded from analysis. 2 donors, N_MP_ = 10, N_Control_ = 6, N_NoCARCtrl_ = 6. Error bars represent mean and standard deviation.

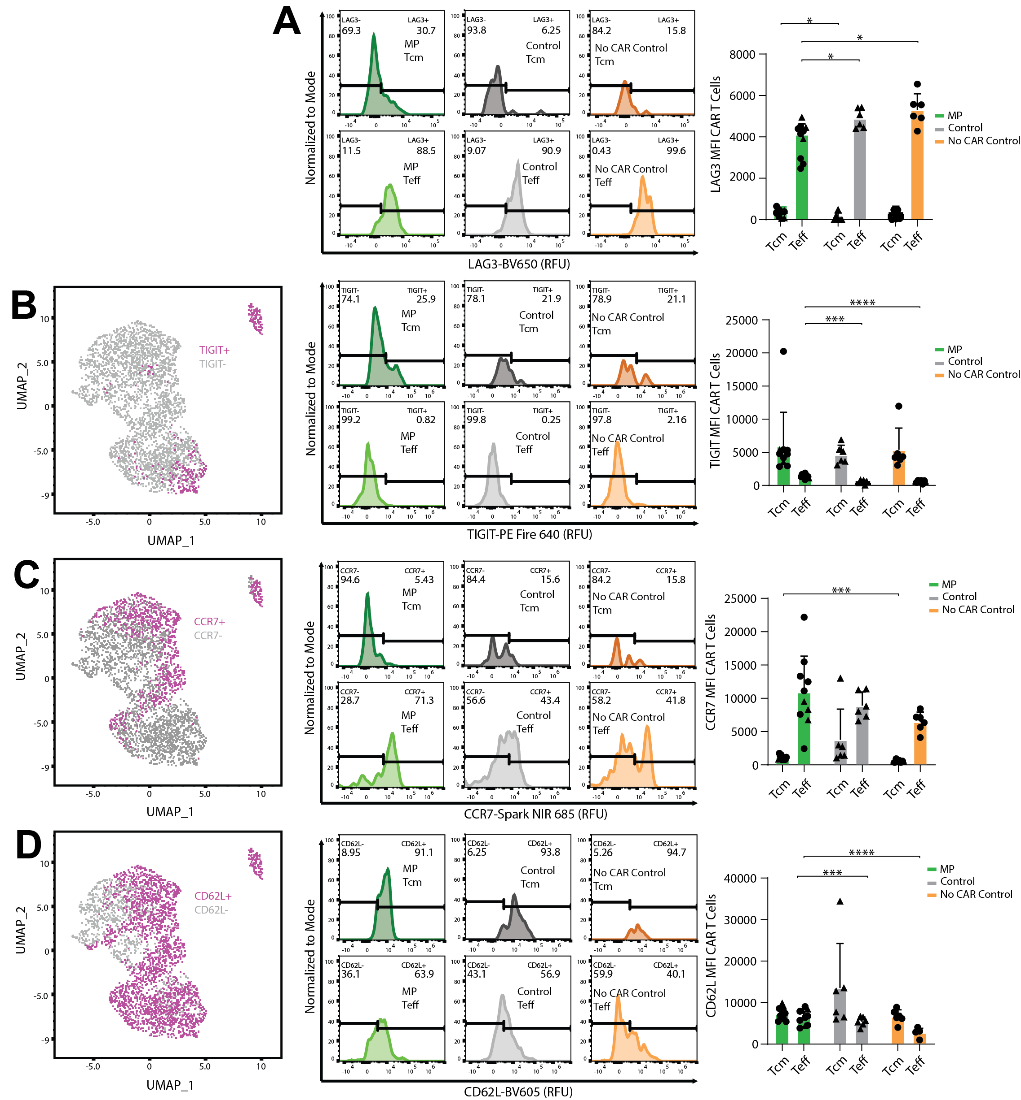

**Figure S9. Expression of Exhaustion and Memory Markers in Splenic Lymphocytes.** The expression of **(A)** LAG3 (Tcm: MP: 730 (342), Control: 115 (200), No CAR Control: 325 (325), p (MP vs Control) = 0.0014) (Teff: MP: 4117 (508), Control: 4895 (467), No CAR Control: 5311 (774), p (MP vs Control and MP vs No CAR Control) = 0.0278 and 0.0274 ), **(B)** TIGIT (Tcm: MP: 5931 (5118), Control: 4654 (1422), No CAR Control: 5381(3282))(Teff: MP: 1407 (288), Control: 521 (287), No CAR Control: 571 (220), p (MP vs Control and MP vs No CAR Control) < 0.001 for both), **(C)** CCR7 (Tcm: MP: 1152 (355), Control: 3784 (4601), No CAR Control: 578 (201), *p* (MP vs No CAR Control) = 0.003) (Teff: MP: 10919 (5421), Control: 8916 (2030), No CAR Control: 6468 (1484)) and **(D)** CD62L (Tcm: MP: 7268 (1437), Control: 13513 (10744), No CAR Control: 6655 (1635)) (Teff: MP: 6535 (1842), Control: 5430 (1122), No CAR Control: 2818 (994), *p* (MP vs No CAR Control) < 0.001, *p* (Control vs No CAR Control) = 0.005) in CD5+/CD45+/TCR-/transgene+ splenic lymphocytes are depicted on UMAP plots, representative histograms and bar graphs for Tcm and Teff. 2 donors, N_MP_ = 10, N_Control_ = 6, N_NoCARCtrl_ = 6. Error bars represent mean and standard deviation. Statistical significance was determined with a Brown-Forsythe and Welsch ANOVA test using Dunnett’s T3 test for multiple comparisons; *p<0.05; ***p<0.001; ****p<0.0001.

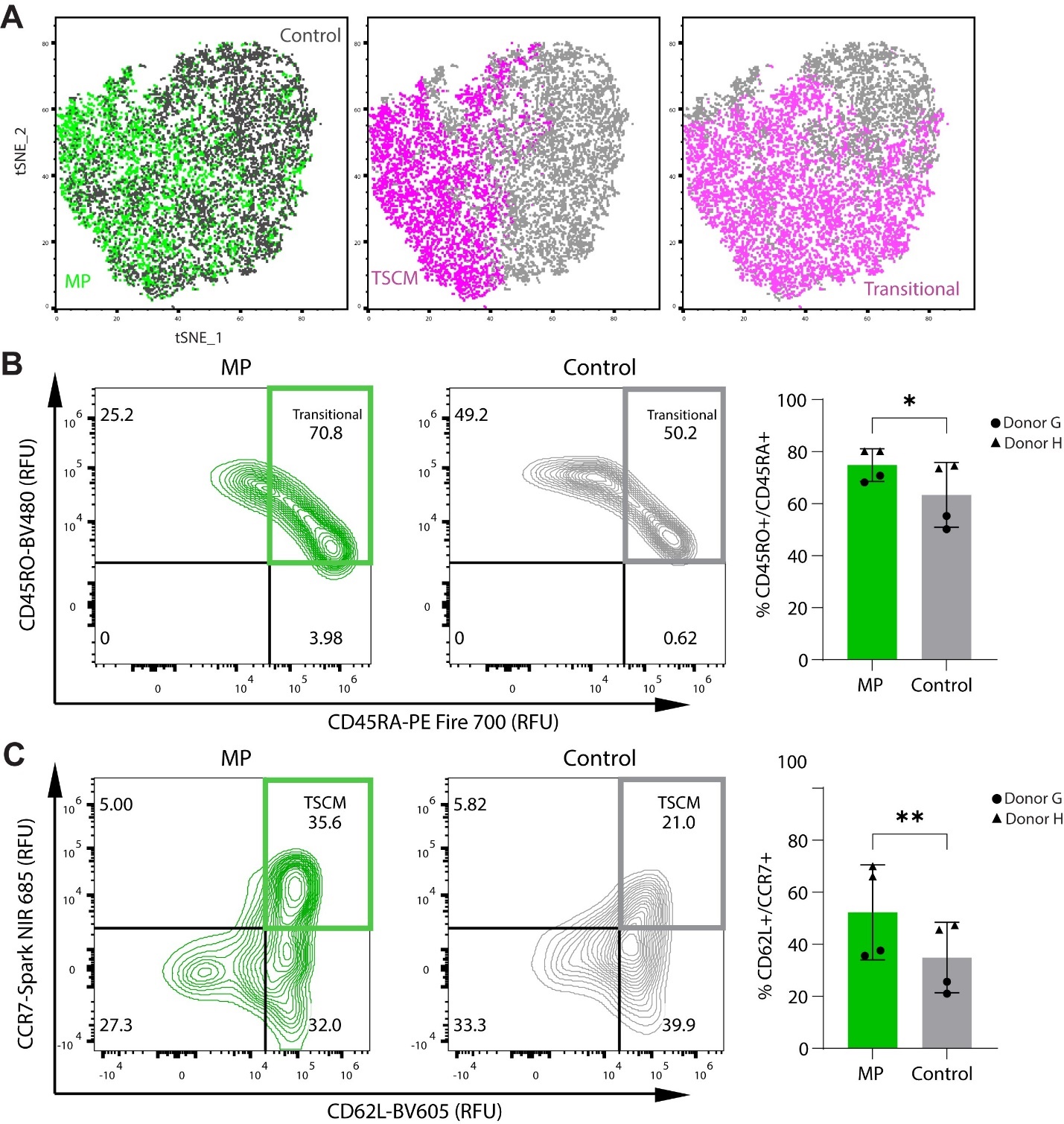

**Figure S10. Differentiation of *TRAC-*CAR T cells in Serial Stimulation Assay.** MP and Control *TRAC-*CAR T cells were serially stimulated with GD2^+^ CHLA-20 neuroblastoma cells for 20 days, collected, and stained to immunophenotype human T lymphocytes via flow cytometry. **(A)** Marker expression t-SNE (t-distributed Stochastic Neighbor Embedding) plots of MP and Control *TRAC-*CAR T cells. These maps were generated via flow cytometry to track CD45RA, CD45RO, CD62L, CCR7, LAG3, and TIGIT expression. Dot plots separate cells by condition, T_SCM_, or transitional T-cell status. Representative contour plot and bar graphs of double positive populations of **(B)** CD45RO vs CD45RA and **(C)** CD62L vs CCR7. 2 donors, N_MP_ = N_Control_ = 4. Error bars represent mean and standard deviation. Statistical significance was determined with paired t-tests; *p<0.05; **p<0.01.

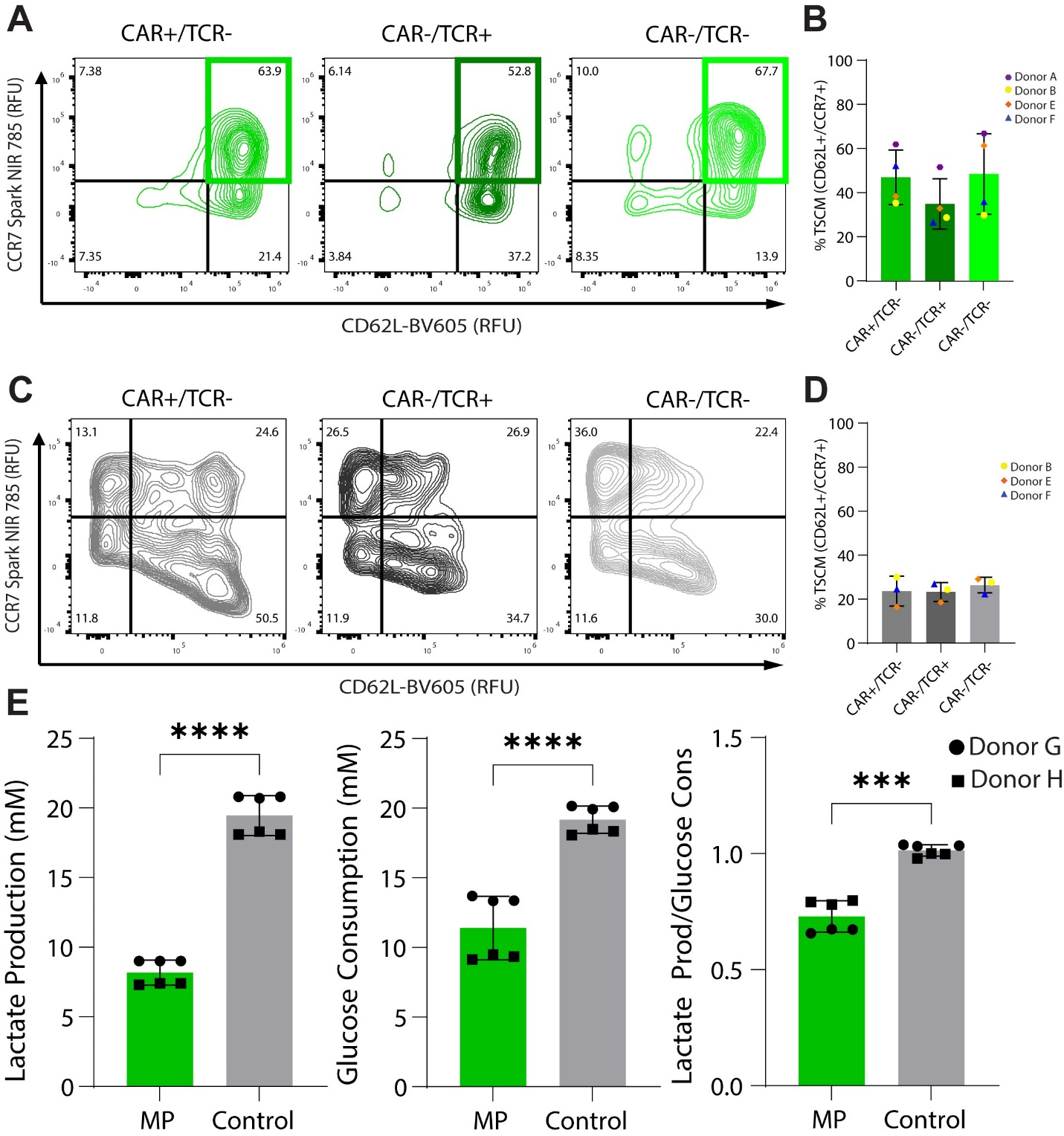

**Figure S11. Expression of Stem Cell Memory Markers in Different T-cell Populations**. Representative contour plots of CD62L vs CCR7 expression and bar graphs of double positive percentages in CAR^+^/TCR^-^, CAR^-^/TCR^+^, or CAR^-^/TCR^-^ populations of **(A,B)** MP (CAR^+^/TCR^-^: 47% (12), CAR^-^/TCR^+^: 35% (11), or CAR^-^/TCR^-^: 48% (18)) or **(C, D)** Control T-cells (CAR^+^/TCR^-^: 24% (7), CAR^-^/TCR^+^: 23% (4) , or CAR^-^/TCR^-^: 26% (4)). **(E)** Untransfected T-cells were grown for 10 days under MP or Control culture conditions where the lactate production (MP: 8.2 mM (0.9), Control: 19.5 mM (1.4), *p* < 0.001), glucose consumption (MP: 11.4 mM (2.3), Control: 19.2 mM (1.0), *p* < 0.001), and the lactate production over glucose consumption (MP: 0.73 (0.07), Control: 1.01 (0.03), *p* < 0.001) was measured on Day 10. Error bars represent mean and standard deviation. Statistical significance was determined with paired t-tests; ***p<0.001; ****p<0.0001.

**Table 1. Guide RNA’s and Primers Used in Study.** The TRAC gRNAs and forward and reverse primers for amplifying the original GD2-CAR and No CAR Control linear constricts for nanoplasmid construction. (5' --> 3')

| **Oligo** | **Sequence** |
| --- | --- |
| TRAC gRNA | CAGGGTTCTGGATATCTGT |
| TRAC PCR REV primer | TAAGGCCGAGACCACCAATCAG |
| TRAC PCR FWD Primer | TCGAGTAAACGGTAGTGCTGGG |

**Table S2. Nanoplasmid Sequences Used in Study.** The full GD2-CAR and No CAR Control nanoplasmid DNA sequences are shown (5' --> 3') (in Excel addendum)

**Table S3. Sanger Sequencing Primers Used in Study.** The primers used for performing Sanger sequencing on the GD2-CAR or No CAR Control plasmid are shown. (5' --> 3') (in Excel addendum)

**Table S4. Antibodies Used in Study for Flow Cytometry.** The clone, manufacturer, fluorophore, catalog number, and volume needed per sample are listed for each antibody used for flow cytometry in this study. (in Excel addendum)
